# Binary vector origin predictably determines *Agrobacterium*-mediated transformation outcome across eukaryotic kingdoms

**DOI:** 10.1101/2025.11.18.688525

**Authors:** Matthew J. Szarzanowicz, Michael Busche, Ziyu Dai, Nathan Lapp, Gina M. Geiselman, Jiayuan Jia, Lucas M. Waldburger, Yan Liang, Guoliang Yuan, Rylan Duong, Kyle R. Pomraning, Nathan J. Hillson, Jacob O. Brunkard, Blake A. Simmons, John M. Gladden, Joonhoon Kim, Mitchell G. Thompson, Patrick M. Shih

**Affiliations:** Joint BioEnergy Institute, 5885 Hollis Street, Emeryville, CA 94608, USA; Environmental Genomics and Systems Biology Division, Lawrence Berkeley National Laboratory, Berkeley, CA, USA; Department of Plant and Microbial Biology, University of California, Berkeley, CA 94720, USA; Laboratory of Genetics, University of Wisconsin-Madison, Madison, WI, USA; Energy and Environment Directorate, Pacific Northwest National Laboratory, 902 Battelle Boulevard, Richland, WA 99354, USA; Sandia National Laboratories, Livermore, CA, USA; Department of Electrical Engineering and Computer Science, University of California, Berkeley, CA 94720, USA; Department of Bioengineering, University of California, Berkeley, California, USA; Biological Systems & Engineering Division, Lawrence Berkeley National Laboratory, Berkeley, CA 94720, USA

## Abstract

*Agrobacterium*-mediated transformation (AMT) is the primary means of genetic engineering in plants and many fungi, but the factors that control transformation outcomes—efficiency, transgene insertion number, and transgene integrity—remain poorly characterized. Although transformation outcomes dictate an event’s potential utility in both industrial and academic contexts, AMT remains largely unoptimized for these metrics. Here, we systematically analyze the impact of the transgene-harboring binary vector on transformation outcomes across plant and fungal species. Through a comparison of different plasmid origin of replication (ORI) families and engineered copy number variants, our results reveal that the ORI family—not plasmid copy number—dictates T-DNA insertion number, backbone inclusion, and transformation efficiency, while plasmid copy number tuning alters efficiency without changing ORI family-specific signatures. Independent of plasmid copy number across kingdoms, the most widely used pVS1 ORI-based vectors (e.g. pCambia) result in significantly more insertions per transformant and high levels of transgene silencing compared to the less-utilized pSa ORI family, which enriches for more uniform single insertion events. Furthermore, we demonstrate that ORI-dependent transformation outcomes in yeast predictably reflect those in *Arabidopsis*. Together, these results lay the foundation for future binary vector design aimed at achieving more predictable, controllable, and optimized transformation outcomes across diverse eukaryotic hosts.

## Introduction

Agrobacterium-mediated transformation (AMT) is the predominant method for introducing transgenes into plants and fungi across academia and industry. Despite its utility, AMT remains an imprecise tool, producing transformants with unpredictable insertion number, concatemerization, and vector backbone inclusion—collectively termed event outcome. These outcomes directly determine a transformant’s suitability for specific engineering goals, and due to the wide range of systems that utilize AMT, ideal event outcomes can vary substantially. For example, single-copy, backbone-free insertions are preferred in conventional crop engineering for regulatory compliance and transgene stability, whereas multi-copy or concatemerized insertions are desirable in metabolic or gene engineering applications to increase enzyme abundance or maximize delivery of homology-directed repair templates. Such diversity of target outcomes makes controllable transformation highly coveted yet largely unachievable with current legacy tools. Thus, unlocking the full potential of AMT will require developing and characterizing components that enable predictable, tunable event outcomes across diverse plant and fungal hosts.

The primary chassis of AMT is derived from the plant pathogen *Agrobacterium tumefaciens* which has a natural capacity to horizontally transfer DNA to plants. A variety of strains and transgene-harboring binary vectors were developed in the 1980s and 1990s, and they continue to form the core toolsets used for AMT today, often fully unchanged from their original design^1^. While AMT generally produces cleaner, lower-copy insertions than sister techniques—e.g., biolistics—reported rates of transgene insertion and integrity vary widely within the literature, making it difficult to determine how each legacy component affects transformation outcome. The sources of variance contributing to these disparate results are numerous and could include host genetic background, host tissue source, *Agrobacterium* strain, transgene architecture, transformation protocol, and the binary vector itself^2–5^.

As the carrier of the transgene, the binary vector plays a major role in influencing transformation outcome. Being a fully synthetic component, each binary vector must contain a user-selected ORI to enable proper replication within *Agrobacterium*. Different ORIs have variable copy number maintenance, stability and partitioning characteristics, potentially changing how the transgene is maintained and processed within the bacterium. Although multiple studies have reported that the binary vector ORI influences transformation outcomes, their conclusions vary—particularly regarding the role of high-copy origins^5–7^. These disparities can likely be attributed to some combination of differences in the plant genetic background, bacterial strain, use of standard vs. super binary vectors, transformation protocol and T-DNA payload. As a result, direct comparisons from the literature are often convoluted for plant transformation due to variations in experimental design.

To systematically assess the impact of binary vector copy number on AMT, we previously constructed a library of 71 copy number variants across four ORIs, each differing by single SNPs in the *repA* gene^8^. These variants were built into otherwise identical binary vectors and evaluated for their influence on transient and stable transformation, demonstrating that both could be significantly improved with single SNPs. Our findings revealed that copy number alone could not explain the observed transformation changes—highlighting that each ORI likely has intrinsic, origin-specific properties that influence transformation. However, a key question remains regarding how plasmid copy number, transformation efficiency, and event outcome are interrelated—factors that collectively define the functional utility of any engineered variant.

To dissect this relationship in a controlled manner across eukaryotic kingdoms, we pursued three primary objectives. First, we characterize the relationship between ORI family, variant-specific copy number, transgene integration number, transgene integrity and backbone inclusion rates using a controlled experimental design in both the model plant *Arabidopsis thaliana* and oleaginous yeast *Rhodosporidium toruloides*, along with validation in the filamentous fungus *Aspergillus niger*. This establishes a foundation for understanding how ORI identity and copy number jointly influence transformation efficiency and outcome. Second, we assess whether GFP expression can serve as a reliable proxy for transgene insertion number in stable transformants, providing a scalable alternative to labor-intensive methods for quantifying insertions. Finally, we test whether transformation outcomes in *R. toruloides* can predict corresponding results in *A. thaliana*, thereby evaluating the potential of using a high-throughput fungal system to predict transformation outcomes in a plant. Our results reveal a key means to improve AMT efficiency while controlling for transformation outcomes across eukaryotic kingdoms. Additionally, we demonstrate a method for high-throughput predictive evaluation of binary vector performance, laying the groundwork for the systematic exploration and engineering of ORI sequence space.

## Results

### ORI family and copy number influence transformation efficiency and GFP expression

As the transgene-harboring plasmid, the copy number of the binary vector within *A. tumefaciens* and the ORI used to control plasmid replication have both been shown to alter stable and transient expression efficiencies^8^. To determine how both the ORI family and plasmid copy number influence transformation outcomes, we selected two ORI families for analysis: pVS1 and pSa. The pVS1 ORI was selected due to its widespread use in the plant community through vectors such as the pCambia series along with its generally high transformation efficiency while pSa was chosen for its low efficiency WT form that had the largest improvement from copy tuning in our previous study^8^. For these families, three variants spanning a low, medium, and high copy number were generated using pVS1 (2, 10, and 29 copies) and pSa (5, 18, and 31 copies) backbones, varying within a family by a single SNP. These vectors were otherwise identical in architecture, with each containing a selectable marker (kanamycin for *A. thaliana*, nourseothricin for *R. toruloides*) near the left border (LB), and a constitutively expressed GFP cassette near the right border (RB). To control for sequence context effects near the borders, randomized 100 bp buffering sequences were added between functional elements and border sequences and standardized across all constructs. These vectors were then introduced into standard *A. tumefaciens* strains for their respective host (GV3101::pMP90 for *A. thaliana* and EHA105 for *R. toruloides*) and used to obtain ∼96 independent transformants per ORI variant, yielding 559 transformants for *A. thaliana* and 576 for *R. toruloides*. These transformants were then used to analyze the relationship between ORI family, vector copy number, population GFP expression, and transgene insertion number following the schematic outlined in **Fig. 1**.

**Fig. 1:**
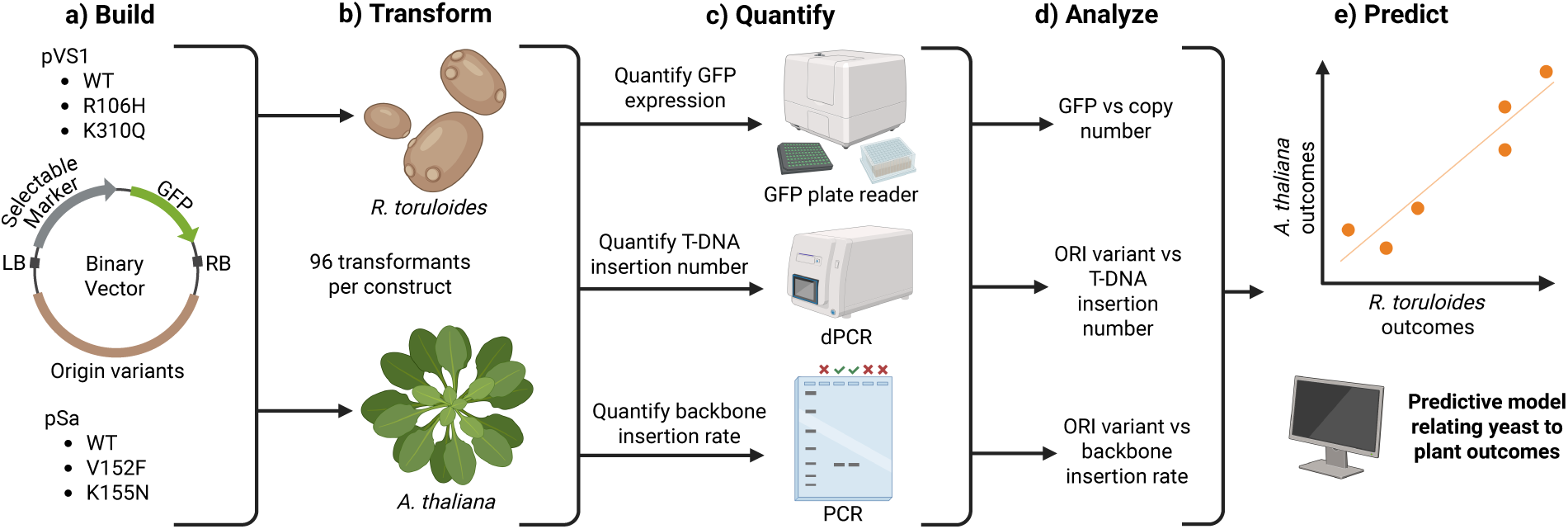
Pipeline analyzing ORI impact on AMT outcomes. **(a)** Six binary vectors were built per organism, consisting of a selectable marker flanking the LB and a constitutive GFP cassette flanking the RB. These binary vectors are maintained in *A. tumefaciens* by one of two ORIs—pVS1 and pSa—which vary in copy number due to single SNPs in the *repA* ORF. **(b)** Strains of *A. tumefaciens* harboring these vectors were used to transform *R. toruloides* and *A. thaliana*, with ∼96 transformants collected per ORI variant. **(c)** These transformants were then analyzed for GFP expression, transgene copy number, and vector backbone inclusion. **(d)** Relationships between GFP expression, transgene copy number, backbone inclusion, and ORI variant were then analyzed, **(e)** and these were used to compare outcome between organisms to evaluate potential predictive capacity between a high-throughput yeast system and a low-throughput plant.

As all transformants for a given species were derived from identical T-DNA sequences harboring a constitutive GFP cassette, variability in GFP expression is likely the product of genomic insertion location, transgene insertion number, transgene silencing, and potentially T-DNA degradation or damage. As AMT-mediated insertions have been well-characterized to occur randomly throughout the genome^9,10^, GFP expression levels across a population would be expected to not differ significantly between binary vector ORI variants if this factor does not influence transformation outcome. Similarly, if differences are observed between ORI variants, it is likely that such variance is due to the impact of the ORI on either transgene insertion number or T-DNA degradation as insertion location and silencing are host-mediated responses. To evaluate potential differences in gene expression, GFP output was measured in both systems using a plate reader assay. For *A. thaliana*, four leaves per transformant were selected, and a single leaf disk per leaf was hole-punched and placed into a 96-well plate for quantification. The disk values were then averaged to produce a final expression value to better account for leaf to leaf variability in expression (Fig. S1a). For *R. toruloides*, liquid cultures were grown in a 96-well block overnight to an OD_600_ between 0.1 and 1.0, and an aliquot of this culture was transferred to a 96-well plate for GFP quantification normalized to OD.

The two selected ORI families vary in their WT copy numbers, with pSa maintaining the binary vector at approximately five copies per cell and pVS1 at ten copies^8^. Through the use of single *repA* SNPs, this number can be manipulated (**Fig. 2a**). Even when comparing variants between ORI families with relatively similar copy numbers, transformation efficiencies tended to vary significantly. This is particularly true for *R. toruloides* (**Fig. 2b**) which typically has more consistent transformation efficiencies between replicates than the highly variable floral dipping of *A. thaliana* (**Fig. 2c**). Transformation efficiencies from *A. thaliana* floral dipping have been well documented to vary by an order of magnitude or more between batches, complicating direct comparisons across variables^11^. For this reason, we used a large sample size of seeds (∼38,000-80,000 seeds per variant) derived from 27-39 independent plants. Due to this high degree of variability across floral dips, a generalized linear model (GLM) was used to estimate transformation efficiencies, accounting for both the seed number and transformation efficiency (**Fig. 2c**). Results from both *R. toruloides* and *A. thaliana* show that the ORI family and sub-variants influence transformation efficiency, corroborating a trend previously seen in^8^. ORI variants with similar estimated copy numbers from different ORI families—e.g. pSa V152F (31 copies) and pVS1 R106H (29 copies)—tended to produce significantly different transformation efficiencies from one another, particularly in *R. toruloides* where this pairing differed by an average efficiency of 16.5-fold across replicates. This provides further evidence that intrinsic ORI family properties are a major source of variance in AMT outcomes, extending beyond copy number alone.

**Fig. 2.**
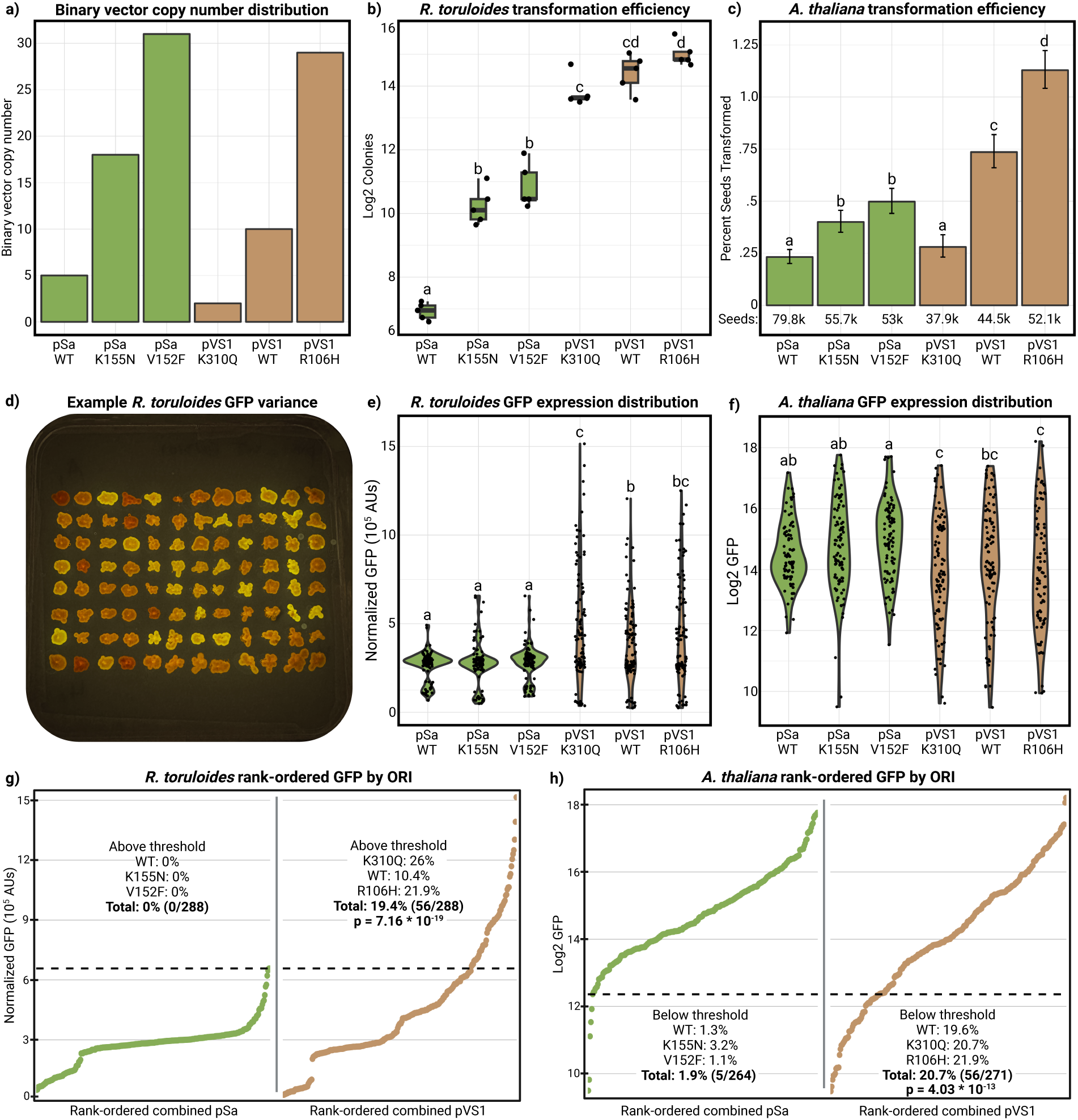
Transformation efficiency and GFP expression vary by ORI family. **(a)** The six ORI variants varied in copy number using *repA* SNPs as characterized in ^8^. Both ORI families and some variants had significant differences in transformation efficiency as measured by total colony number for *R. toruloides* **(b)** and percent transformed seed for *A. thaliana* **(c)**. Significant differences were determined by a two-tailed ANOVA followed by a Tukey’s post hoc test for *R. toruloides* (p < 0.05 for significance letters) and a binomial GLM with estimated marginal means and Wald 95% CIs for *A. thaliana*. Transformants from the same transformation varied dramatically in GFP expression **(d)**, and this variance was found to be primarily ORI family—not vector copy number—dependent for both *R. toruloides* **(e)** and *A. thaliana* **(f)**. Population level expression differences were determined by a two-tailed ANOVA followed by a Tukey’s HSD test (p < 0.05 for significance letters). When rank-ordering the samples by GFP expression, it was observed that the pVS1 ORI family had significantly more transformants above the pSa-population maximum (19.4%, p = 7.16 * 10^-16^) for *R. toruloides* **(g)** but significantly fewer than the lower boundary (2nd percentile) of the continuous pSa family range for *A. thaliana* **(h)** (20.7%, p = 4.03 * 10^-13^) as measured by a hypergeometric test.

When analyzing population-level GFP outputs, ORI-dependent differences became even more pronounced. Across both systems, GFP expression distributions were highly consistent within each ORI family, despite differences observed on the copy number and transformation efficiency levels. Within the *R. toruloides* distributions, transformants derived from any pSa variant exhibited lower average and less variable GFP expression than pVS1-derived transformants. Contrastingly, pVS1-derived samples displayed a broader range of expression, with a long upper tail extending toward highly and visually-distinct GFP expressing colonies (**Fig. 2e and 2d**). The results in *A. thaliana* were intriguingly the opposite: pVS1-derived lines were found to have a long lower tail comprising transformants with little to effectively no GFP expression (**Fig. 2f**). When rank-ordering all samples by GFP output, 19.4% (56/288) of *R. toruloides* pVS1-derived transformants exceeded the maximum pSa expression value across all pSa variants (n = 288, **Fig. 2g**). Conversely, 20.7% (56/271) of pVS1-derived lines in *A. thaliana* were found to fall below the general pSa population’s minimum, with only 1.9% (5/264) of pSa outliers doing the same (**Fig. 2h**). In both species, around 20% of the pVS1-derived samples notably deviated beyond the extremes of the pSa population. This species-dependent expression variability may reflect gene silencing in *A. thaliana* that is triggered upon hyper-expression of a transgene^12^—a process that may be absent or attenuated in some fungi compared to plant systems.

To further explore this observation, a filamentous fungal system, *Aspergillus niger*, was transformed with the same six ORI variants, and 96 transformants were evaluated per vector. These parallel experiments reproduced the family-level patterns with pVS1 variants yielding markedly higher transformation efficiencies and a broader, right-shifted GFP distribution compared to pSa (Fig. S2, b–c). Thus, across all systems, the ORI family was observed to exhibit significant impacts on both transformation efficiency and transgene expression, and notably, these expression trends remained even when controlling for plasmid copy number across ORI families.

### Transgene insertion number is ORI family-dependent and influences GFP expression

The differences observed in GFP expression between ORI families are likely due to different transgene insertion numbers across the population. To determine if the ORI family and copy-tuned variants altered transformation outcome, gDNA was extracted from all samples and prepared for transgene insertion quantification via dPCR. For each transformant, two probes targeting the GFP cassette and a genomic target were used within the same reaction, and the transgene insertion number was estimated from the ratio of GFP to genomic copies, adjusting for ploidy. dPCR enables absolute quantification of starting template copies, offering greater accuracy and precision than qPCR and higher-throughput multiplexing compared to Southern blotting, making it well-suited for large-scale transgene studies^13^.

Similar to the GFP data, notable ORI-dependent differences in insertion number distributions were observed for both organisms (**Fig. 3a and 3b**). All pSa ORI variants produced significantly lower average insertion numbers and higher rates of single insertions than pVS1 binary vectors across systems (63% single insertions for combined pSa samples in *R. toruloides* vs 40% for pVS1, and 36% vs 13% in *A. thaliana*). A notable difference in the total number of insertions was observed between organisms, with 95% of transformants having fewer than six insertions in *R. toruloides* compared to 35 copies in *A. thaliana* (**Fig. 3c** and **Fig. 3d**). Of particular note, all pVS1 vectors in *A. thaliana* produced high-insertion transformants that reached an estimated 140 insertions, coupled with low rates of single-insertion events. Such high insertion numbers are likely attributable to concatemerization of T-DNAs rather than exclusive insertion into independent loci. Transformation of germline tissues, such as floral dipping, is known to produce higher rates of T-DNA concatemerization compared to somatic transformation, and large concatemerized T-DNA insertions have been previously reported in floral dipped *A. thaliana*^14–16^.

**Fig. 3:**
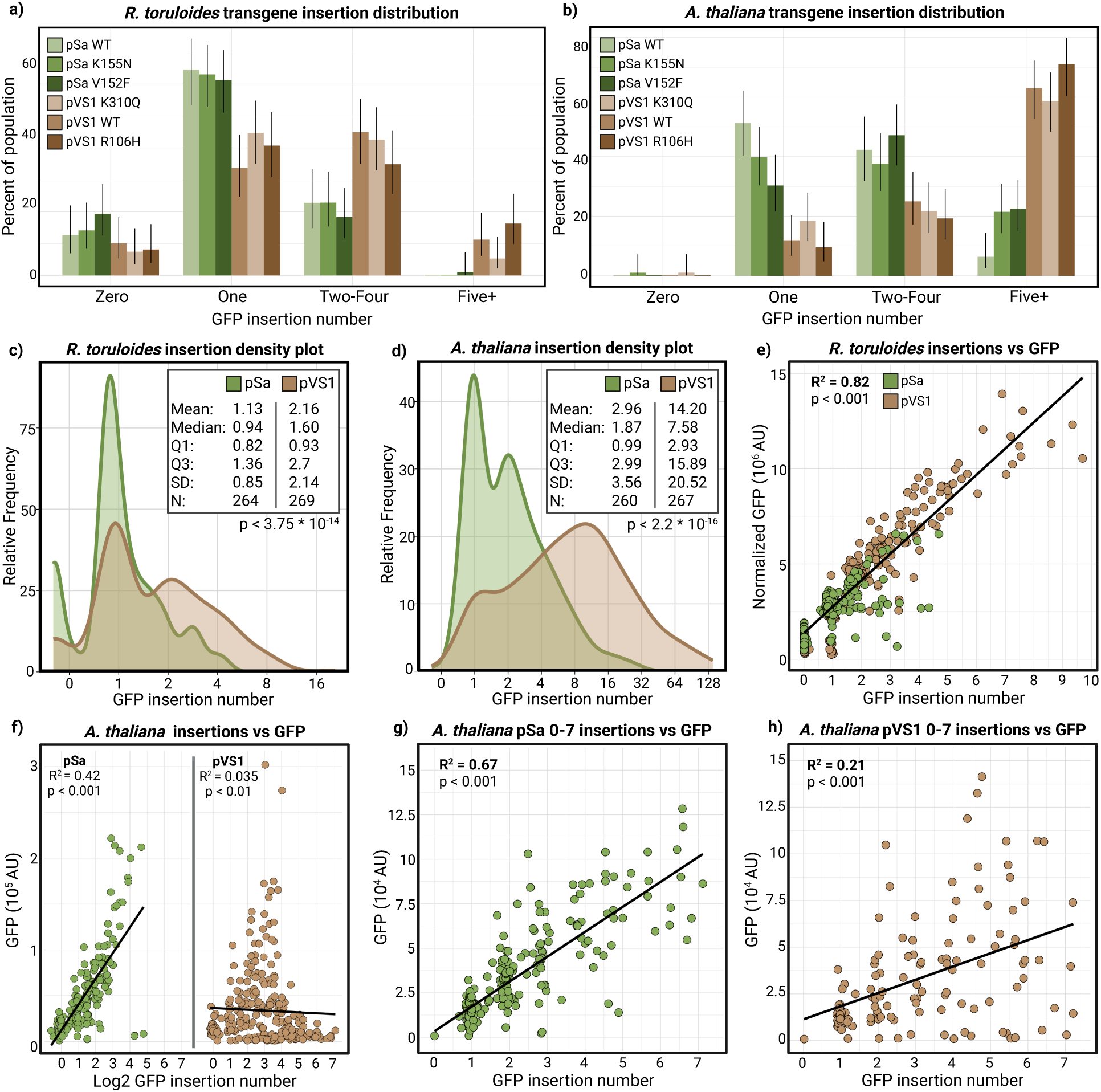
Transgene insertion number and GFP expression vary by ORI family and organism. Transgene insertion numbers were estimated with dPCR and grouped by 0, 1, 2-4, and 5+ insertions for *R. toruloides* **(a)** and *A. thaliana* **(b)**. Population percentage averages for each species were calculated from the estimated marginal means of a binomial GLM along with Wald 95% CIs, depicted as error bars. The ORI family significantly altered the rate of transgene insertions, with pSa producing both lower average insertion numbers and higher rates of single insertions than pVS1 for *R. toruloides* **(c)** (p < 3.75 * 10^-14^) and *A. thaliana* **(d)** (p < 2.2 * 10^-16^) as computed by a Wilcoxon test. GFP followed a linear gene dosage regression in *R. toruloides* **(e)** (R^2^ = 0.82, p < 2.2 * 10^-16^ from the F statistic) that fit both ORI families. **(f)** In *A. thaliana*, the pVS1-derived population had a substantial proportion of low-expressing high-insertion transformants, reducing the variance explained by insertion number to effectively zero (adjusted R^2^ = 0.03, p = 2.04 * 10^-3^) while the pSa-derived population retained some of the linear relationship (R^2^ = 0.42, p < 2.2*10^-16^). When looking only at transformants with fewer than 8 insertions in *A. thaliana*, the pSa **(g)** and pVS1 **(h)** populations differed significantly in the linear relationship between GFP and insertion number despite being transformed with the same T-DNA sequence (pSa: adjusted R^2^ = 0.67, p < 2.2 * 10^-16^ and pVS1: adjusted R^2^ = 0.21, p = 3.61*10^-8^).

Such high insertion numbers were not observed in *R. toruloides* for either ORI, nor for the pSa variants in *A. thaliana*, making this an exclusive feature of pVS1 vectors in *A. thaliana*, regardless of plasmid copy number. Moreover, as the pVS1 ORI family is broadly used in plant transformation within vector series such as pCambia and pPZP, this finding is notable and cautionary for plant transformation. As further validation of higher insertion rates for pVS1, we quantified GFP transgene insertion number in *A. niger* transformants derived from WT pSa (n = 93) and pVS1 vectors (n = 92). Consistent with the other systems, pVS1-derived transformants had significantly higher average insertion numbers than pSa (**Fig. S2 d–e**, p < 0.001), further corroborating this ORI-family level trend.

Within *A. thaliana* and *R. toruloides*, all pVS1 copy number variants produced single insertions at non-significantly different rates (**Fig. S3**, p > 0.05). A similar lack of significant intra-family differences was observed among pSa variants in both systems, although in *A. thaliana* the high-copy pSa V152F variant trended toward a reduction in single-insertion rate when compared to pSa WT (**Fig. S3**, p = 0.069). Taken together, these results demonstrate that plasmid copy number, transformation efficiency, and single-insertion generation rates are not inextricably linked, suggesting possible future engineering to improve efficiency without compromising single-insertion event generation.

To evaluate the relationship between gene dosage and expression, estimated GFP insertion numbers were plotted against GFP expression for each transformant. For *R. toruloides*, a strong linear relationship was observed between insertion number and expression, indicating a predictable gene dosage effect. A Cook’s D-adjusted linear regression yielded an R² of 0.82, indicating that insertion number accounts for the majority of expression variance, with minimal influence from positional effects or silencing (**Fig. 3e, Fig. S4).** Both pSa- and pVS1-derived transformants aligned along this trendline, suggesting minimal influence of ORI family on transgene expression outcomes within *R. toruloides*.

In contrast, *A. thaliana* exhibited a far more complex pattern with pronounced ORI-dependent effects on transgene expression. Transformants with insertion numbers exceeding 16 almost exclusively exhibited low but detectable GFP expression, likely indicating transgene silencing which is a well documented phenomenon in plants. Across all insertion number ranges, there was at least a small proportion of individuals with similarly low levels of expression caused either by silencing, transgene degradation, or positional effects within the genome (e.g. heterochromatic insertion). Despite the >200-fold variance in expression levels, a general relationship between gene dosage and expression could be defined at lower insertion numbers, declining after seven insertions as silencing became more prevalent. Among transformants with seven or fewer insertions, a weaker gene dosage relationship was observed (R² = 0.43) relative to *R. toruloides*, demonstrating greater variability attributable to positional effects or silencing in the plant system. Intriguingly, splitting the data by ORI family increased the predictive capacity dramatically for pSa (R² = 0.67, n = 243) but decreased it for pVS1 (R² = 0.21, n = 132) (**Fig. 3g and 3h**). This difference can largely be attributed to a greater number of low expressing—likely silenced—transformants derived from pVS1 binary vectors. As both the pSa and pVS1 vectors carried identical T-DNA sequences across all variants, these differences are ORI-dependent, demonstrating that the binary vector ORI can profoundly influence not only transgene insertion rate but also downstream expression levels within *A. thaliana*. Why identical T-DNA sequences launched from different binary vector backbones yield variable expression outcomes for a given insertion number remains unknown; however, one could speculate that pVS1-launched T-DNA undergoes greater rates of concatemerization or degradation compared to pSa, leading to changes in host silencing or transgene integrity.

Taken as a whole, these results show that the binary vector ORI family—not the vector copy number—profoundly impacts transgene insertion rates across systems. Within *A. thaliana*, the ORI family exhibits an additional effect of influencing transgene expression for a given insertion number, making the selection of the binary vector ORI important not just for controlling insertion rates but also transgene expression quality.

### ORI family influences backbone insertion but not T-DNA integrity in *A. thaliana*

The impact of the ORI family on transgene expression suggests this feature of the binary vector may influence transgene mobilization or integrity. To further investigate how the ORI family changes transformation outcome, we evaluated T-DNA integrity and backbone insertion frequencies within the *A. thaliana* transformant population (**Fig. 4a**). T-DNA degradation has been widely reported within the literature and results in truncated insertions that do not span the entire RB-LB sequence^17,18^. As mobilized T-DNAs are covalently bound to VirD2 at their 5’ RB end, there is a general assumption within the community that LB degradation is more common due to susceptibility to exonuclease activity^3^. As such, it is common practice to place the selectable marker near the LB to help purge LB-degraded transformants during selection.

**Fig. 4:**
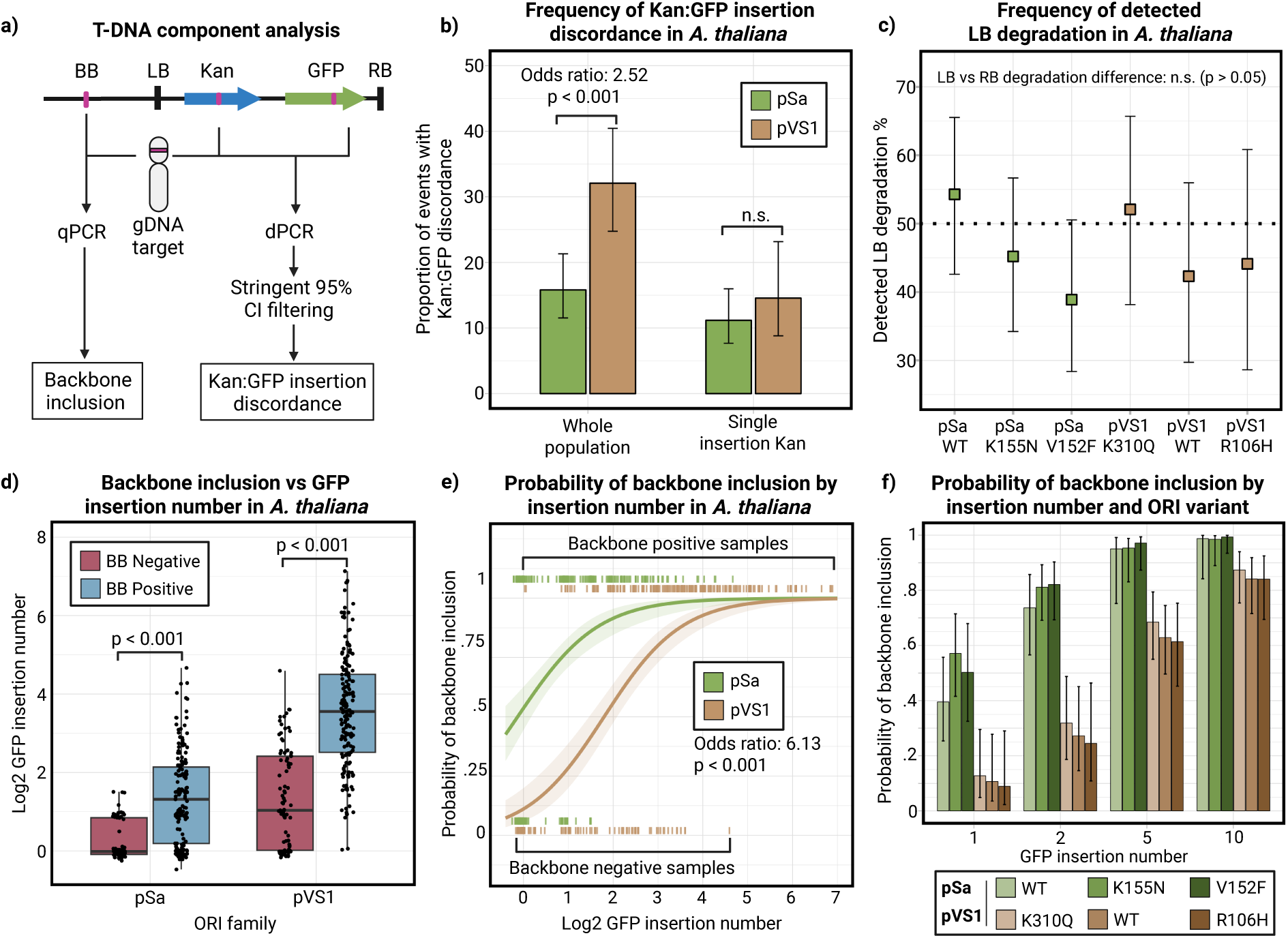
Evaluation of T-DNA integrity and backbone inclusion in *A. thaliana.* **a)** Schematic of the T-DNA integrity and backbone inclusion analysis. T-DNA integrity was assessed by dPCR of the LB-proximal *Kan* and RB-proximal *GFP* cassettes, each normalized to a gDNA control. Backbone inclusion was quantified by qPCR of a locus ∼300 bp from the LB and normalized to the same gDNA control. For integrity analyses, only samples with 95% CI ranges from the dPCR Poisson quantification confined to a single integer copy for both *Kan* and *GFP* were retained. **b)** Incidence of discordant *Kan:GFP* ratios was modeled with a binomial GLM. The predicted probabilities of discordance from the estimated marginal mean of the GLM + 95% Wald CI are shown. Across the full population, pVS1 transformants (n = 134) showed higher discordance than pSa (n = 215; odds ratio = 2.52, *p* = 0.00045). In single-insertion lines, rates did not differ (pSa n = 101, pVS1 n = 30; *p* > 0.05). **c)** Border truncations were inferred from *Kan:GFP* imbalance and analyzed by ORI variant using a binomial GLM. Predicted probabilities from emmeans were compared against the null expectation of equal LB and RB degradation (50%) via Wald tests. No variant (pSa n = 70–73, pVS1 n = 34–52) deviated significantly (*p* > 0.05). **d)** Backbone inclusion vs. log2 GFP insertion number compared by a Wilcoxon rank sum test for pSa (n = 243, p = 2.35 * 10^-12^) and pVS1 (n = 267, p = 1.37 * 10^-23^). **e)** A GLM including ORI family, GFP insertion number, and backbone inclusion was used to test family-level effects. Wald tests of emmeans contrasts revealed that pSa transformants (n = 243) had a significantly greater odds of backbone inclusion per GFP insertion than pVS1 (n = 267; odds ratio = 6.13, *p* = 2.15 * 10^-12^). **f)** Predicted probabilities of backbone inclusion at defined GFP insertion numbers were estimated for each ORI variant using emmeans from the GLM; error bars represent 95% Wald CIs.

We investigated LB and RB border degradation by quantifying insertion numbers of the kanamycin resistance (Kan) and GFP cassettes, positioned near the LB and RB, respectively. Using the same DNA templates as the GFP analysis, we performed additional dPCR assays to determine Kan insertion number, with probes targeting Kan and a genomic reference. Each probe provides an absolute template count with an associated 95% confidence interval based on Poisson partitioning. Thus, for each sample we obtained four independent error estimates (GFP, Kan, and two genomic controls). To reduce false positives, we only considered transformants where both targets had 95% CIs confined within a single integer insertion number. After filtering, 215 transformants were evaluable for pSa and 134 for pVS1, with no event exceeding 12 insertions of either cassette due to the higher quantification error associated with elevated insertion number.

When comparing the insertion number of Kan to GFP within this high confidence population, we found significantly higher rates of Kan:GFP insertion number discordance within pVS1-derived transformants than pSa-derived transformants (odds ratio 2.52x, p = 0.00045), suggesting higher rates of T-DNA degradation within the pVS1 population (**Fig. 4b**). When considering only events with single insertions of Kan, discordance was similar between the two ORI families (n.s., p > 0.05), implicating the higher average insertion rate of all pVS1-variants to the higher discordance on the population level. As this analysis only measures the ratio of the two genes, the rates of detected RB or LB degradation are likely underestimated due to masking in multi-insertion transformants with equal numbers of LB and RB degraded T-DNAs.

For each transformant with an unequal Kan:GFP insertion ratio, an estimate of LB or RB degradation can be made based on the surplus of Kan or GFP, indicating RB and LB degradations or truncations, respectively. To test the LB-degradation hypothesis from exonuclease activity, we analyzed the rate of LB degradation using a generalized linear model across ORI variants. This analysis found no enrichment of LB degradation for any ORI variant (p > 0.05), suggesting incomplete T-DNA insertions occur at similar rates from both the RB and LB borders and that ORI families and sub-variants do not significantly influence this property (**Fig. 4c**). These unanticipated results challenge the notion of greater LB degradation and suggest the common design practice of localizing the selectable marker to the LB deserves reconsideration.

In addition to T-DNA degradation, numerous reports have characterized the inclusion of the binary vector backbone (BB) in transformants. This phenomenon is most commonly observed as a LB readthrough, in which termination at the LB sequence does not properly occur^19^. Within the literature, backbone inclusion rates vary widely, typically ranging between 20-80% of transformants^7,20–24^. The wide variability in this reported range likely reflects variation in transformed species, *Agrobacterium* strain, binary vector architecture, transformation protocol and sampling size, making it difficult to identify significant sources of variance related to backbone mobilization.

To analyze the impact of the binary vector ORI on backbone inclusion, we quantified inclusion rates for all *A. thaliana* transformants using qPCR for a backbone target ∼300bp from the LB along with a gDNA internal control. Transformants cleanly separated into backbone-positive and negative groups when comparing the Cq values for the backbone and gDNA targets (**Fig. S5a**). When comparing backbone inclusion and GFP insertion number, both ORI families showed significant enrichment of transgene insertions in backbone positive lines, suggesting the probability of backbone inclusion increases in multi-insertion lines (**Fig. 4d**, p < 0.001 for both pSa and pVS1). Despite the differences in average insertion numbers between ORI families, however, backbone inclusion rates for all ORI variants did not significantly differ and ranged from 58-79% (**Fig. S5b,** p > 0.05 with Holm-corrected GLM). As a transformant needs only one backbone insertion to be classified as backbone-positive, it is reasonable to hypothesize that transformants with higher total insertion numbers (due to more independent T-DNA insertions or concatemerization) also have a higher chance of having at least one included backbone. To investigate this, we implemented a generalized linear model incorporating backbone inclusion, ORI family, and measured GFP insertions to analyze the probability of backbone inclusion as a function of GFP insertion number. This analysis found that the pSa ORI family had a significantly higher chance of backbone inclusion per copy of GFP than pVS1 (**Fig. 4e**, odds ratio: 6.13x, p < 0.001). As pSa variants tend to produce transformants with notably lower insertion numbers than the pVS1 family, this difference is masked on the population level as abundant high-insertion pVS1-derived transformants tend to have backbone.

When comparing backbone inclusion rate probabilities for each ORI variant at different GFP insertion numbers, variants from each ORI family clustered tightly and were not significantly different (**Fig. 4f**). This suggests that the differences observed in backbone inclusion are an ORI family-dependent property and are not significantly impacted by manipulations in plasmid copy number that affect other properties such as transformation efficiency. Why certain ORI families influence the rate of backbone inclusion on otherwise identical binary vectors is unknown. There is literature precedence of certain ORIs producing lower rates of backbone inclusion such as the pRi ORI compared to RK2^7^, but to date, there is no validated mechanistic explanation to describe this phenomenon. This study provides strong evidence that such differences are derived from intrinsic properties of the ORI itself in a manner that is independent of binary vector copy number.

### GFP expression is a functional predictor of transgene insertion number

All core analyses in this study relied upon large-population transformation, gDNA extraction, and dPCR-based quantification. This pipeline is highly labor intensive, costly, and not scalable to screening thousands of ORI variant candidates, even within most industrial settings. To address these limitations in throughput, we investigated whether GFP expression could serve as a reliable proxy for predicting the number of inserted transgenes. Given the broad and irregular distribution of expression levels observed in the *A. thaliana* dataset, we implemented an ensemble of machine learning models to better capture nonlinear trends and improve performance across a variably expressing population. Models were trained using either GFP expression alone or GFP combined with ORI identity to assess the added predictive value of the ORI family.

To enhance resolution, transformants were grouped into three biologically relevant categories based on transgene insertion number: single insertions, 2–4 insertions, and five or more insertions. A similar strategy was applied to *R. toruloides*, with the addition of a zero-copy group to account for transformants with no detectable GFP insertions despite viable growth on nourseothricin.

For both species, we initially evaluated a diverse set of individual classification models (**Fig. S6**). From these, the top three performers—each utilizing distinct classification strategies—were selected to form a majority-vote ensemble in which predictions for unknown samples are determined by consensus. In *A. thaliana*, the selected models were Support Vector Classification (SVC), Naive Bayes (NB), and Quadratic Discriminant Analysis. For *R. toruloides*, the ensemble consisted of SVC, NB, and a multinomial logistic regression model.

Each model was evaluated under both GFP-only and GFP+ORI conditions using a stratified 70:30 training-to-testing split, repeated over 100 bootstrap iterations to estimate average model performance. Classification outcomes are summarized in the confusion matrices shown in **Fig. 5b-c and S7**. These matrices provide the basis for calculating precision and recall metrics—enabling quantification of false positive and negative rates—which were combined to yield the F1 scores shown in **Fig. 5d and S7.**

**Fig. 5:**
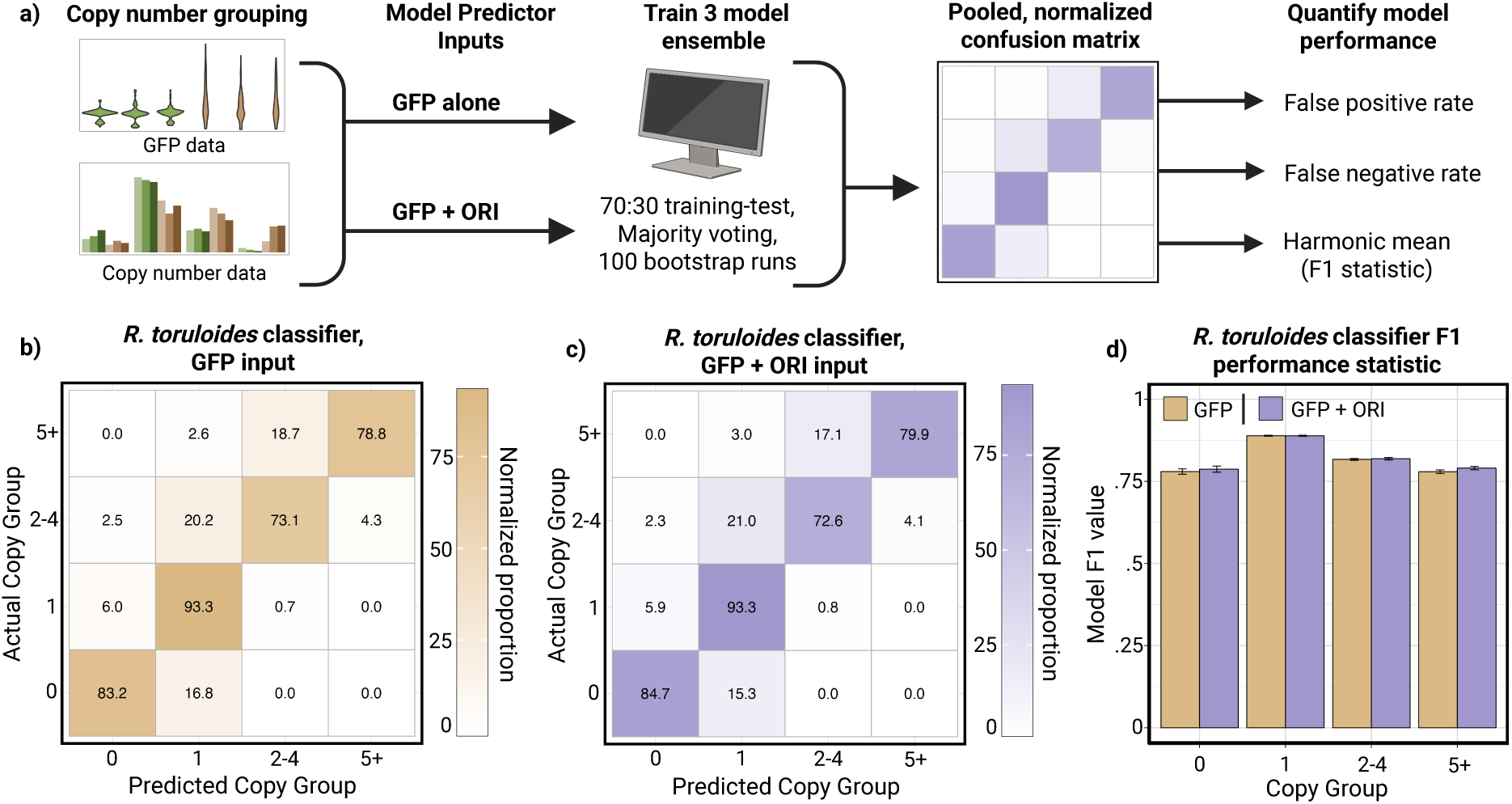
Ensemble prediction of GFP insertion number in *R. toruloides.* **(a)** An ensemble of predictive models was trained to predict the GFP insertion number of a transformant based on either GFP output alone or GFP + ORI identity as predictive factors. These data were used to conduct 100 bootstrapped 70:30 training-test splits across three independent models, which were then combined in a majority voting ensemble. Confusion matrices were generated from the bootstrapped runs to analyze the ensemble’s performance and rates of false positive and negative results. *R. toruloides* showed high predictive capacity of insertion number from GFP alone **(b)** that was not improved by the addition of ORI identity **(c)**. *R. toruloides* ensemble F1 statistics are reported in **(d)**, showing predictive capacity for all groups from GFP alone (F1: 0 insertions, 0.78; 1 insertion, 0.89; 2-4 insertions, 0.82; 5+ insertions, 0.78). Error bars represent the SEM from the bootstrapped runs.

The results of these ensembles show that *R. toruloides* insertion number can be predicted reliably from GFP alone (Mean F1 value of 0.82), with single insertions in particular achieving high predictability (F1 = 0.89). Importantly, including the ORI family did not substantially improve model performance, supporting the notion that the insertion number-GFP trend is ORI-independent in this species, enabling prediction without *a priori* knowledge about an ORI family.

Prediction in *A. thaliana* was more challenging due to greater complexity between GFP expression, insertion number, and silencing. Within this system, GFP expression alone enabled modest classification of single-insertion and five+ insertion transformants (F1 scores = 0.61 and 0.49, respectively) but showed no ability to identify the intermediate 2-4 insertion group (F = 0.01). Including the ORI family identity in the training model substantially improved predictive performance, increasing F1 scores to 0.75, 0.62, and 0.75 for the 3 groups (**Fig. S7**). This further supports the previous observation of ORI-dependent expression outcomes within *A. thaliana*.

In summary, GFP expression can serve as an excellent proxy for transgene insertion number in *R. toruloides*, providing a promising avenue for high-throughput insertion number estimation using simple fluorescence measurements. In *A. thaliana*, GFP alone provides some, but more limited, information about insertion number extremes (single vs five+ insertions), and inclusion of ORI identity is needed to achieve finer resolution. As it is possible that ORIs beyond the two used in this study will perform differently cased on the variance observed, *a priori* information about the ORI will be needed to train ORI-specific models in *A. thaliana* to maximize performance in this species.

### Intra- and interspecies modeling using GFP for copy number prediction

Having shown that GFP is a viable proxy for insertion number in *R. toruloides*, we next evaluated the model’s external validity and generalization. To do this, we took the same six ORI variants and conducted another transformation, selecting 96 transformants per variant for evaluation. In the same manner as before, these transformants were grown overnight and measured for GFP fluorescence, which was then used for ensemble prediction of the insertion number group. These predicted population-level insertional distributions were compared to their measured reference populations to determine if there was significant similarity with previous results. For all ORI variants across all insertion groups, predicted class proportions fell within the 95% posterior-predicted confidence intervals of the measured insertion number classes (Dirichlet–multinomial), and there were no significant differences between the measured and predicted population outcomes (**Fig. 6a**, Fisher’s Exact Test with Monte-Carlo, FDR-adjusted; all q > 0.05). This demonstrates that GFP-based modeling recapitulates *R. toruloides* population compositions in new transformations without gDNA/dPCR, validating fluorescent measurements and ensemble prediction as an accurate alternative to manual quantification.

**Fig. 6.**
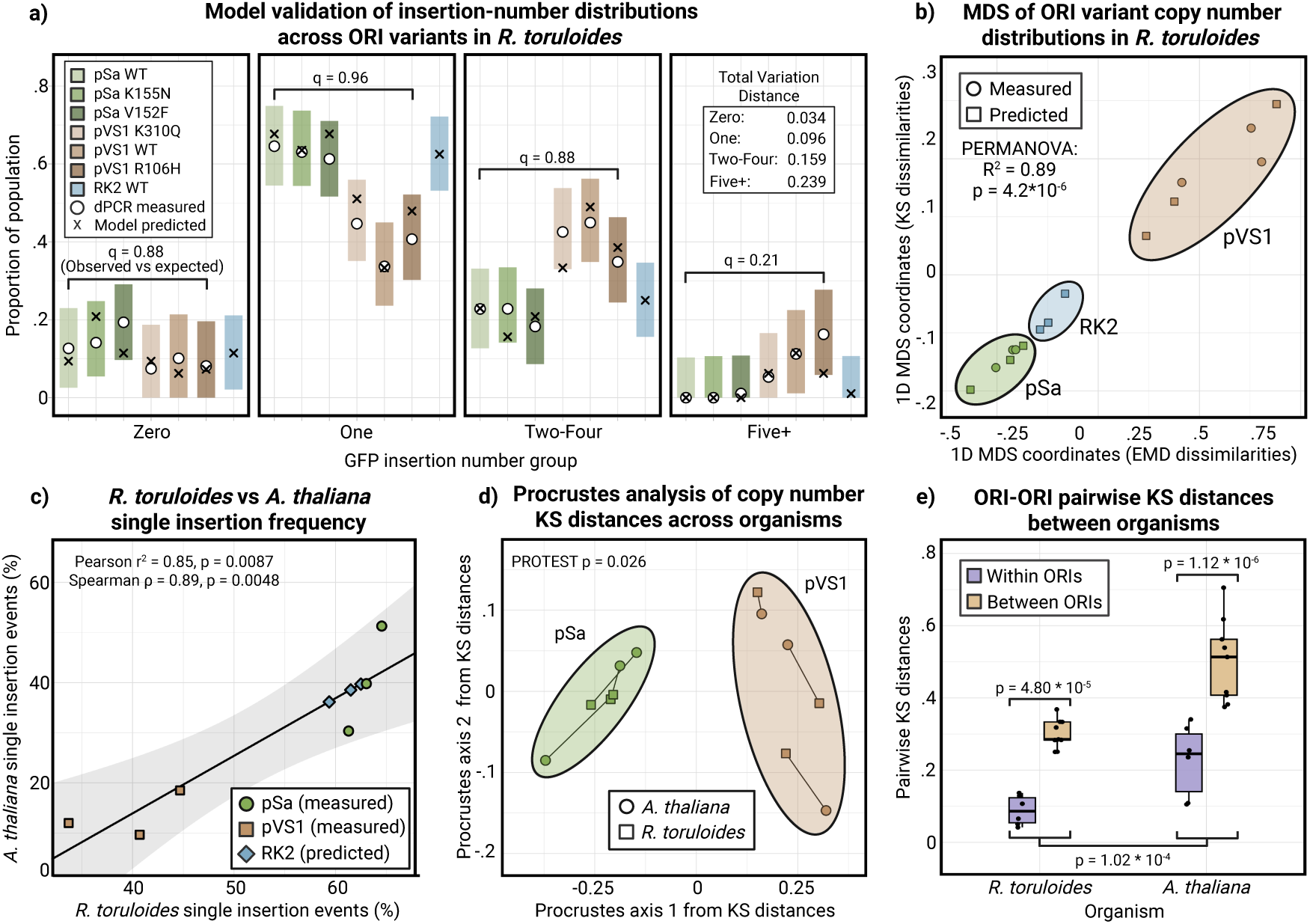
Intra- and inter-organism modeling of transformation outcomes. **(a)** *R. toruloides* model validation by ORI variant. For each variant, bars show the measured population proportion for each GFP insertion-number class with 95% simultaneous multinomial intervals (Sison–Glaz). Markers denote the measured (o) or new population predicted (x) proportion for each group. Each group was evaluated with a Fisher’s Exact Test followed by Monte-Carlo resampling (FDR adjusted; all q > 0.05). The inset reports the total-variation distance (TV) between measured and predicted outcomes. **(b)** Multidimensional scaling (MDS) of ORI-variant copy number distributions using 1D embeddings from Kolmogorov–Smirnov (KS) and Earth Mover’s Distance (EMD) dissimilarities. Points show measured and predicted populations for pSa and pVS1 variants; RK2 is shown for predicted only. Ellipses indicate significant ORI-family clusters (R^2^ = 0.89, p = 4.2*10^-6^). **c)** Linear regression of *A. thaliana* versus *R. toruloides* single GFP insertion frequencies across the six shared ORI variants; the shaded band is the 95% CI of the fit, derived from the standard errors of the slope and intercept. (Pearson r^2^ = 0.85, p = 0.0087; Spearman ρ = 0.89, p = 0.0048). RK2 predicted points are shown for reference but excluded from the regression. **(d)** Procrustes analysis comparing the pairwise KS-distances of ORI variants between organisms (measured data only). Lines connect matched ORI variants between organisms and ellipses denote ORI families. A Procrustes Randomization Test (PROTEST) indicates significant congruence of the two configurations (p = 0.026). **(e)** ORI–ORI pairwise KS distances within versus between ORI families in each organism. Boxplots summarize the distribution of within-family and between-family distances; overlaid p-values are from ANOVA contrasts. The between-within ORI variant trends are concordant across organisms and do not differ significantly (see **Fig. S8**).

Having validated the reproducibility of model results on ORI variants that were used to train the ensemble, we next evaluated ensemble performance on a new unseen ORI family, RK2. Using the WT form (∼1 copy per cell) and two *repA* copy number variants—S20F (∼5 copies) and R11G (∼15 copies)–-we transformed *R. toruloides* with binary vectors containing identical T-DNA cargo as previous experiments. Measuring the GFP expression from 96 transformants per ORI variant, we estimated the population GFP insertion distribution using the ensemble. We then embedded all populations—the original pSa and pVS1 measured transformants, their prediction-only replicates, and novel RK2 data—via multi-dimensional scaling on pairwise 1D KS and EMD distances (**Fig. 6b**). Measured and prediction-only populations co-localized, and all three ORI families separated significantly (Permutational ANOVA, p = 4.2 * 10^-6^), confirming that fluorescence-based predictions recapitulate ORI-family level clustering. All RK2 variants clustered more closely to the pSa family than pVS1 variants, strongly suggesting that RK2 binary vectors produce transformation outcomes that are more similar to pSa than the pVS1 family, with greater numbers of single insertions and lower average insertion number. Together, these results provide a proof-of-concept pipeline in which new ORIs can be rapidly screened from fluorescence in ∼1 week in *R. toruloides* without the need for costly and laborious gDNA extraction and transgene quantification for exploratory analyses.

With the *R. toruloides* modeling validated, we next asked whether the population-level signals in *R. toruloides* were reflected in *A. thaliana*. To explore this, we analyzed the measured single-insertion event generation frequency for all ORI variants in both systems. Single insertion events are a valued transformation outcome for most plant engineering, making this a metric of particular importance. Plotting the population outcomes against one another found a strong and significant correlation between the two systems (**Fig. 6d**, Pearson r^2^ = 0.85, p = 0.0087; Spearman ρ = 0.89, p = 0.0048).

To expand the analysis beyond single-insertion transformants, we summarized whole-population differences by MDS of pairwise KS distances among ORIs for each organism (**Fig. S8**). These species-level MDS configurations were then compared using a Procrustes analysis which found significant alignment between ORI variants across species (**Fig. 6e**, PROTEST p = 0.026). This analysis implies that inter-ORI outcomes are conserved across organisms, despite possible differences in overall effect size. To further explore how population variance between all pairwise ORI-ORI groupings correlated between organisms, pairwise KS distances were calculated and plotted against one another, resulting in 15 total comparisons per species. This comparison showed significant concordance of ORI-ORI variant differences across organisms, particularly for differentiating ORI-family level differences (**Fig. S9**, p < 1.0 * 10^-6^). We further divided these pairwise comparisons into within-ORI-family and between-family groups and found consistent differences between organisms with between-family comparisons having greater KS distances than within-family pairings (**Fig. 6f**, p < 0.001). On the species level, *A. thaliana* was found to have a generally higher insertion rate and greater variability for all ORI variants than transformants in *R. toruloides*. This is reflected in both the insertion spread and larger KS distance magnitudes, but comparing within- and between-family distances showed no significant differences between organisms, further suggesting a general cross-species trend (**Fig. S9**).

Taken as a whole, this modeling suggests two core results: 1) GFP can accurately predict transformation outcomes in *R. toruloides*, and 2) population data in *R. toruloides* has a significant predictive capacity on transformation outcomes in *A. thaliana* on both the single-insertion level and ORI-ORI variance level more broadly. This pipeline of modeling GFP fluorescence to estimate population insertional distributions in *R. toruloides* followed by transformation outcome prediction in *A. thaliana* opens the door to streamlined screening of ORI variants, bypassing bottlenecks that typically limit plant-based systems.

## Discussion

In this study, we systematically evaluated how the binary vector ORI and plasmid copy number influence AMT outcomes across eukaryotic kingdoms. Our results demonstrate that ORI family—not copy number—is the primary determinant of transformation efficiency, transgene insertion number, transgene expression, and vector backbone inclusion frequency. ORI-dependent differences persisted even when plasmid copy number was substantially altered, revealing intrinsic, ORI-specific properties that strongly shape transformation outcomes.

While binary vector copy number was observed to influence transformation efficiency, its impact on transgene insertion frequency was dramatically lower compared to the influence of the underlying ORI family. These findings are particularly important given legitimate concerns that engineering ORIs to improve transformation efficiency could compromise quality by increasing transgene insertion number and backbone inclusion rates. In this study, we did not observe consistent or substantial tradeoffs between plasmid copy number and transformation quality across the variants and organisms tested. While modest effects were detectable—for example the pSa V152F variant in *Arabidopsis*—such impacts were not a general feature of the dataset. As such, there appears to be an engineerable space where single-insertion rates or transformation efficiency can be improved without substantially compromising other transformation variables, although broader testing across additional ORI families and transformation contexts will be necessary to fully validate this approach.

Since similar plasmid copy number variants across different ORI families produced highly variable results, intrinsic ORI characteristics appear to be major determinants of AMT outcomes. Our findings suggest that these ORI-specific features—such as potential differences in T-DNA supercoiling, plasmid localization within *Agrobacterium*, or interaction with host factors—retain their influence even when plasmid copy number is extensively modified. Notably, disparities in transgene expression in *A. thaliana* among transformants with identical insertion numbers underscore an underappreciated role for the binary vector ORI’s influence on transformant quality. These results carry significant implications for users of pCambia vectors, which are widely used in academia for plant transformation but rely on the pVS1 ORI. In *A. thaliana*, pVS1-based vectors consistently produced high multi-insertion events—reaching up to 140 copies per transformant—and were associated with elevated rates of transgene silencing. These outcomes raise concerns about the reliability of pCambia-derived transformants in *Arabidopsis* research. Given these findings, we recommend reevaluating the use of pCambia vectors in *A. thaliana* and conducting a more systematic assessment of alternative ORIs for this system. For any engineering effort that would benefit from higher proportions of single insertions and functional transformation efficiency, utilizing the pSa K155N variant in *A. thaliana* and pSa K155N or V152F for *R. toruloides* are recommended from the variants tested in this study. For high insertion rates across systems, the pVS1 R106H vector can be deployed to combine high efficiency transformation with high rates of insertion.

Given the notable effects observed at the ORI family level, expanding bioprospecting efforts to explore the diverse natural repertoire of ORIs is a critical next step for improving AMT. To date, conducting such a screen and evaluating transformation outcomes directly in plants—even efficient models like *Arabidopsis*—remains prohibitively low-throughput. To address this bottleneck, we developed a high-throughput fungal proxy system using *R. toruloides* coupled with GFP fluorescence as a predictive indicator of transformation outcome. Comparison of results between this yeast system and *A. thaliana* revealed significant mirrored trends across organisms for both rates of single insertions and general ORI-ORI variant differences. This proxy system represents a promising advance to enable scalable, predictive evaluation of binary vector performance without the throughput limitations imposed by direct plant assays. Future research should prioritize validating the generalizability of this system across a wider range of ORIs, host organisms, and vector architectures. By implementing a high-throughput proxy system, this approach has the potential to accelerate the exploration of ORI sequence space and advance AMT toward becoming a more predictive and controllable platform for transgene insertion.

## Methods

### Media, chemicals and culture conditions

Bacterial cultures were grown in LB Miller medium (BD Biosciences) shaking at 200 rpm at 37 °C for *E. coli* and 30 °C for *A. tumefaciens* unless otherwise noted. Cultures were supplemented with kanamycin (50 mg L^-1^; Sigma-Aldrich), gentamicin (30 mg L^-1^; Thermo Fisher Scientific), or rifampicin (100 mg L^-1^; Teknova) as indicated. All compounds unless otherwise noted were purchased from Sigma-Aldrich.

### Strains and plasmids

All bacterial strains and plasmids used in this study are listed in Supplementary Table 1 and are viewable through the public instance of the Joint BioEnergy Institute (JBEI) registry (https://public-registry.jbei.org/folders/931). All strains and plasmids created in this work can be requested from the strain archivist at JBEI with a signed material transfer agreement. All plasmids were Gibson assembled using standard protocols^25^ or Golden Gate assembled using standard protocols^26^. Plasmids were routinely isolated using the QIAprep spin miniprep kit (Qiagen), and all primers were purchased from Integrated DNA Technologies (IDT). Plasmid sequences were verified using whole-plasmid sequencing (Plasmidsaurus). *Agrobacterium* was routinely transformed by electroporation as described previously using a 1-mm cuvette and a 2.4-kV, 25-μF, 200-Ω pulse^27^.

### Arabidopsis thaliana transformation

The Columbia-0 (Col-0) ecotype of *Arabidopsis thaliana* was grown in Sunshine no. 4 medium supplemented with Osmocote at 22C under long-day conditions (16 h light, 8 h darkness; 150 μmol m^−2^ s^−1^ PAR) for 2 weeks after sowing numerous seeds onto the surface of a pot.

Seedlings were then transplanted to 4-inch pots, with 3 seedlings per pot, and grown under the same conditions until flowering. The primary floral inflorescence was removed after reaching ∼2 cm to encourage axillary shooting. When the inflorescences had grown and set their first few siliques, the pots of plants were transformed via floral dip as previously described^28^.

Briefly, the binary vectors used in this study were transformed into *A. tumefaciens* GV3101 (pMP90), which was grown to saturation in LB medium + kanamycin, gentamicin and rifampicin, shaking at 200 rpm at 30 °C overnight. 100 mL of culture was centrifuged at 3,200g for 15 minutes and resuspended in 200 mL of floral dip medium, consisting of 5% sucrose and 0.02% Silwet. All formed siliques were removed with scissors, and then all inflorescences per pot were submerged into this solution for 30 seconds. For buds that could not reach the solution, a pipette was used to eject 50 μL of bacterial solution onto the bud surface. Up to 4 pots that had been dipped into the same solution were then placed into a tray sideways and covered with a plastic dome that had been misted with water. The following day, the pots were uncovered, returned to an upright position, and placed into a growth chamber with identical growth conditions as above. One week later, this process was repeated without removing developing siliques. Following the second dip, the plants were allowed to continue growing with standard watering for 2 weeks followed by cessation of watering to allow for full desiccation.

### *A. thaliana* selection and evaluation

Fully desiccated plants were harvested by sifting seeds through progressively finer sieves to remove chaff. As seed weight was used to estimate total seed number, care was taken to remove even fine chaff through >20 progressive sifts. ∼150 mg of seed—corresponding to roughly 7500 seeds^29^—were placed into 1.5 mL Eppendorf tubes and sterilized. First, 1 mL of 70% ethanol was placed into each tube for 1 minute, inverting. The ethanol was removed, and 1 mL of 50% commercial bleach (12.5% hypochlorite) + 1 drop of Triton-20 surfactant was added, and the tube was inverted for 7.5 minutes. This sterilization solution was then removed, and the seeds were rinsed 3 times with sterile water. Seeds were then evenly spread onto the surface of 6.5-inch plates containing Murashige and Skoog salts + 1 g L^−1^ MES brought to a pH of 5.7 along with 50 mg L^−1^ kanamycin and 300 mg L^−1^ timentin to kill residual *A. tumefaciens* and 8 g L^−1^ agar for solidification. These plates were placed into growth chambers with identical conditions used for *A. thaliana* and were allowed to grow for 3 weeks.

Following germination and kanamycin selection, 96 transgenic *A. thaliana* were transplanted per construct into flats—1 plant per cell, 24 cells per tray, 4 trays per construct. Plants were allowed to grow under identical long-day conditions for 3 weeks. Following this in a systematic plant-by-plant manner, 4 leaves were removed per plant, and a single 6 mm hole punch was taken per leaf. These leaf discs were placed abaxial side up in a 96-well culture plate containing 330 μL water per well. The remaining tissue was then placed into a separate 96-well block for storage and later genomic DNA extraction. Each 96-well plate containing the leaf discs was then measured for GFP fluorescence using a BioTek Synergy H1 microplate reader set for conditions of 483-nm excitation and 512-nm emission. Raw disc-level *A. thaliana* meta data can be found in Supplementary Table 2. Genomic DNA from each sample was extracted using a 96-sample DNeasy kit (Qiagen, Product 69181).

### *Rhodosporidium toruloides* transformation and evaluation

The binary vectors used in this study were transformed into *A. tumefaciens* EHA105 (pTiBo542 ΔT-DNA). Cultures of *A. tumefaciens* were grown overnight in LB + kanamycin and rifampicin at 30 °C shaking at 200 rpm. Cultures were suspended at an OD_600_ of 1.0 in an AB salts induction medium for 24 hours ^30^. On the same day, *R. toruloides* that had been grown to saturation in YPD medium (BD Biosciences) at 30 °C shaking at 200 rpm was diluted into YPD, such that the culture would reach an OD_600_ between 0.5-1.0 the following morning.

After the induction period, 1 mL of OD_600_ = 1.0 *A. tumefaciens* and 1 mL of OD_600_ = 1.0 *R. toruloides* were mixed and centrifuged at 4000g for 5 minutes. The supernatant was removed, and the pellet resuspended in 1 mL of induction medium. This resuspension was then vacuumed using a central laboratory vacuum line through a Millipore 0.45-micron membrane, which was then placed onto induction medium that had been solidified with 2% agar. The cells were allowed to co-cultivate for 4 days in the dark at 26 °C. Following this period, the pellet was resuspended by vortexing the filter in 5 mL of YPD, and cells were plated on YPD + 2% agar with 100 mg L^-1^ nourseothricin and 300 mg L^-1^ timentin. Selected colonies emerged after 2 days of incubation at 30 °C, and these were quantified using an AnalytikJena UVP GelSolo for imaging and Fiji for counting (https://imagej.net/software/fiji/downloads).

For each construct, 96 colonies were then randomly selected and used to inoculate a 96-well block (500 μL YPD + 20 mg L^-1^ nourseothricin + 300 mg L^-1^ timentin per well) for overnight growth at 30 °C shaking at 200 rpm. The following day, 150 μL of culture was added to a 96-well plate along with 150 μL of YPD, and each plate was measured for both OD_600_ and GFP fluorescence using a BioTek Synergy H1 microplate reader set for conditions of 483-nm excitation and 512-nm emission. Final GFP fluorescence was normalized to culture concentration by dividing measured fluorescence by culture OD_600_. Data for this experiment can be found in Supplementary Table 3. Genomic DNA for all cultures was extracted from the pellet of 1 mL culture grown to an OD_600_ of ∼1.0 using a Quick-DNA Fungal/Bacterial 96-sample kit (Zymo Research, Product D6006).

### *Aspergillus niger* transformation and evaluation

The six binary vectors used in this study were transformed into the EHA105 strain of *A. tumefaciens* by electroporation. Fresh spores of the *albAD* mutant of *Aspergillus niger* derived from the ATCC 11414 stock were produced on *Aspergillus* Complete Medium (CM) and diluted in 0.4% Tween-80 to a concentration of 10^5^-10^6^ spores per 125 mL. This mutant was used to enable better GFP quantification by removing melanin biosynthesis. The *Agrobacterium* cultures were grown in induction medium ^31^ to an OD_600_ of 0.6-1.0 and were then mixed in a 1:1 ratio with the spore solution. This mixture was spread onto Amersham TM hybond-N nylon membranes and placed on induction medium plates, followed by incubation for 2 days at 20 °C. The membranes were then transferred to minimal medium agar plates containing 150 μg/mL hygromycin B and 250 μg/mL cefotaxime and were incubated at 30 °C for 3-4 days.

Transformed colonies were randomly picked and transferred to 13×100 mm sterile glass tubes containing a 1.5 mL CM agar slant with the proper antibiotics. After 3-4 days of growth at 30 °C, spores were released via vortexing and pipetting with 1 mL sterile 0.4% Tween-80 and passed through one-layer Miracloth (EMD Millipore Corp, Burlington, MA, USA) into 1.5 mL microcentrifuge tubes. Spore concentrations were determined by ORFLO Moxi V cell analyzer (ORFLO Technologies, El Cajon, CA, USA), and the GFP fluorescence of each spore culture was measured with a SpectraMax M5 (Molecular Devices, San Jose, CA, USA) using 395 nm excitation and 509 nm emission spectra. GFP fluorescence was normalized to 10^6^ spores for each strain. Data can be found in Supplementary Table 3.

For genomic DNA extraction, spores of each transformant were inoculated into 1.5 mL CM liquid medium with the proper antibiotics in 13 x 100 mm culture tubes and incubated without shaking at 30 °C for 30-40 hours. Biomass was then removed and transferred into 1.5 mL tubes and lyophilized followed by homogenization with glass beads using a mini-beadbeater-16 (BioSpec Products, Inc, Bartlesville, OK, USA). DNA was purified using the Maxwell® RSC plant DNA kit by the Maxwell® RSC instrument (Promega, Madison, WI, USA).

### Transgene insertion number quantification

Transgene insertion number was quantified for all systems using nanoplate digital PCR with the QIAcuity One system (Qiagen, Product 911001). Genomic DNA concentrations were quantified using a Qubit fluorometric system (ThermoFisher Scientific, Product Q33238) and were diluted to a concentration of 10,000-100,000 genomic copies per μL, followed by digestion with organism-specific restriction enzymes that are specified below. All primary quantifications targeted the GFP cassette within the T-DNA and a genomic control target using the primer-probe pairings specified below:

### *A. thaliana* (Digested with NciI)

- GFP (F: TTGTACTCCAGCTTGTGC, R: GACGGCAACTACAAGACC, FAM probe: TTCAGCTCGATGCGGTTCACCAGG)
- gDNA (F: CCATCTACGGCATCAATATCC, R: AAACATTCCCAAGCAGTCC, HEX probe: ACCACTTCCAAGAACATCTGCCACA)
- Kanamycin cassette (additional *A. thaliana* target) (F: TACCGTAAAGCACGAGGA, R: GGACCGCTATCAGGACATA, FAM probe: TCGCCGCCAAGCTCTTCAGCAATA).

### *R. toruloides* (Digested with SacI)

- GFP (F: GGTGTTCTGCTGGTAGT, R: CAGAAGAACGGCATCAAG, FAM Probe: CCTCGATGTTGTGGCGGATCTTGA)
- gDNA (F: CCAGTCACCAACGGGAACG, R: CGACTCCGCTCACGAAGC, HEX probe: AGCGCTATGTCGACGAGCGCTTCCCGTA)

### *A. niger* (Digested with AluI)

- GFP (F: CAGGAACGCACGATCTTC, R: CGATACGGTTAACCAGTGTATC, FAM probe: ACTATAAGACGCGGGCGGAGGTGA)
- gDNA (F: GTACACCCGCAACTTTACC, R: GCAGAGTGCCATTGTATGT, HEX probe: TGGAGGCATTTGACAGCCATGCAGT)

All samples were run in 96-well 8,500-partition QIAcuity nanoplates (Product ID: 250021) using the QIAcuity Probe PCR Mastermix (Product ID: 250102) with 40 2-step cycles of 15 seconds denaturing at 95 °C and 30 seconds annealing and extension at 60 °C. Each reaction contained primers and probes for both the T-DNA target (FAM probe) and gDNA target (HEX probe), enabling absolute quantification of template numbers for both targets in each sample using Poisson statistical methods via the QIAcuity Software Suite. The transgene insertion number was calculated as the ratio of T-DNA:gDNA templates, adjusting for ploidy (*A. thaliana* diploid, fungal systems haploid). Data can be found in Supplementary Table 3.

### Backbone analysis

The inclusion of the binary vector backbone within *A. thaliana* transformants was analyzed using qPCR on a CFX96 (Bio-Rad, discontinued) using SsoAdvanced™ Universal SYBR® Green Supermix (Bio-Rad, Product ID: 1725271). A sequence 40 bp from the left border was targeted with the following primers: F: AAATCACCACTCGATACAGG, R: ACAAGACGAACTCCAATTCA, and the same primer set as the dPCR gDNA target was used for the gDNA control. Each target was run in a separate reaction using 40 2-part cycles consisting of 15 seconds of 95 °C for denaturing and 30 seconds of 60 °C for annealing and extension. Backbone inclusion was defined as any sample with a Backbone Cq – Control Cq that was ≥2, which is indicative of samples with notably lower backbone templates compared to gDNA control target templates. Backbone positive and negative samples separated into distinct groups using this threshold as shown in Fig. S5a. Backbone signal found in all samples was attributed to lysed residual *Agrobacterium* harboring binary vectors that contained the backbone template sequence.

### Insertion number modeling

All modeling analyses were conducted in R using the packages caret, MASS, e1071, randomForest, class, nnet, and glmnet. Data from the combined transformation dataset were filtered by organism, rounded to the nearest integer GFP insertion number, and grouped into three or four categorical classes. For *R. toruloides*, insertions were classified into “Zero,” “One,” “Two–Four,” and “Five+” insertion groups, while *A. thaliana* was divided into “One,” “Two–Four,” and “Five+,” as zero-insertion events were virtually absent in this system. Two predictor sets were evaluated for each organism: one using GFP fluorescence alone, and another combining fluorescence with binary vector ORI family identity.

Eight supervised learning algorithms were benchmarked: quadratic and linear discriminant analysis, k-nearest neighbors (k = 4), random forest, multinomial logistic regression, linear support vector machine (SVM), naïve Bayes, and penalized multinomial regression. Models were trained and tested over 100 iterations of stratified Monte Carlo cross-validation, in which 70% of data were randomly assigned to training and 30% to testing using caret::createDataPartition. For each iteration, model predictions on held-out data were compared to true class labels using caret::confusionMatrix, from which overall and per-class balanced accuracies were extracted. Numeric predictors were scaled where appropriate, and factor levels were standardized across partitions. Mean per-class and overall accuracies were computed across iterations, with the standard error of the mean reported for overall accuracy.

From these benchmark results, a three-model ensemble was constructed to improve prediction robustness. The ensemble combined a linear SVM, naïve Bayes, and multinomial logistic regression classifier for *R. toruloides*, and a linear SVM, naïve Bayes, and quadratic discriminant analysis model for *A. thaliana*. Each model was trained independently using GFP fluorescence (± ORI family identity) as input features, and predictions were aggregated through majority voting, where each model cast one vote for a predicted insertion class. When all three models agreed, the classification was unanimous; when two agreed, the majority class was selected. In the rare case of a three-way tie (each model predicting a different class), the sample was assigned to the “Zero” category to ensure deterministic resolution. This ensemble framework leveraged the complementary strengths of discriminative and probabilistic learners, producing more stable and accurate predictions of GFP insertion number from fluorescence data.

### Statistical Analyses

All statistical analysis was conducted in R using the following packages: core parametric and nonparametric tests—including *t*, Wilcoxon, Kolmogorov–Smirnov, Shapiro–Wilk, and correlation tests—were conducted with the stats package. The emmeans package provided estimated marginal means and pairwise contrasts for generalized linear models. Distributional differences were quantified using Earth Mover’s Distance from emdist, and rstatix was used for pairwise Wilcoxon tests with Benjamini–Hochberg correction. Linear and logistic model summaries were tidied using broom, and multcompView generated compact letter displays summarizing multiple comparisons. For machine learning–based classification, the caret framework was used to benchmark eight algorithms spanning MASS (LDA/QDA), e1071 (Naïve Bayes, SVM), randomForest, class (kNN), and glmnet (penalized regression), with MLmetrics providing F1 score evaluation.

## Supporting information

Plasmids and Strains

## Acknowledgements

We would like to thank Stanton Gelvin and N. Louise Glass for productive discussions related to this manuscript. Figure cartoon representations sourced from www.biorender.com. M.B. and J.O.B. were supported by NIH R01-GM145814. L.M.W. was funded through the National Science Foundation Graduate Research Fellowship Program. *Aspergillus* work was performed by the Pacific Northwest National Laboratory, operated for the U. S. Department of Energy (DOE) by Battelle under Contract DE-AC06-76RL01830 and supported by the U.S. DOE’s Bioenergy Technologies Office under agreement 2.5.3.106. Other work was part of the JBEI (https://www.jbei.org) supported by the US Department of Energy (DOE), Office of Science, Office of Biological and Environmental Research, through contract DE-AC02-05CH11231 between Lawrence Berkeley National Laboratory and the DOE. The funders had no role in manuscript preparation or the decision to publish. The views and opinions of the authors expressed herein do not necessarily state or reflect those of the United States government or any agency thereof. Neither the United States government nor any agency thereof, nor any of their employees, makes any warranty, expressed or implied, assumes any legal liability or responsibility for the accuracy, completeness or usefulness of any information, apparatus, product or process disclosed or represents that its use would not infringe privately owned rights. The United States government retains and the publisher, by accepting the article for publication, acknowledges that the United States government retains a nonexclusive, paid-up, irrevocable, worldwide license to publish or reproduce the published form of this manuscript or allow others to do so for United States government purposes. The DOE provides public access to these results of federally sponsored research in accordance with the DOE public access plan (http://energy.gov/downloads/doe-public-access-plan).

## Contributions

Conceptualization, M.J.S., M.G.T., P.M.S.; methodology, M.J.S., M.B., Z.D., N.L., G.M.G., J.J., M.G.T; investigation, M.J.S., M.B., Z.D., J.J., L.M.W., Y.L., M.G.T., P.M.S.; writing—original draft, M.J.S.; writing—review and editing, all authors; resources and supervision, N.J.H., J.O.B., B.S., J.M.G., J.K., M.G.T., P.M.S.

## Bibliography

1. Hellens, R., Mullineaux, P. & Klee, H. Technical Focus:a guide to Agrobacterium binary Ti vectors. Trends Plant Sci. 5, 446–451 (2000).

2. Chetty, V. J. et al. Evaluation of four Agrobacterium tumefaciens strains for the genetic transformation of tomato (Solanum lycopersicum L.) cultivar Micro-Tom. Plant Cell Rep. 32, 239–247 (2013).

3. Gelvin, S. B. Integration of Agrobacterium T-DNA into the Plant Genome. Annu. Rev. Genet. 51, 195–217 (2017).

4. Grevelding, C., Fantes, V., Kemper, E., Schell, J. & Masterson, R. Single-copy T-DNA insertions in Arabidopsis are the predominant form of integration in root-derived transgenics, whereas multiple insertions are found in leaf discs. Plant Mol. Biol. 23, 847–860 (1993).

5. Zhi, L. et al. Effect of Agrobacterium strain and plasmid copy number on transformation frequency, event quality and usable event quality in an elite maize cultivar. Plant Cell Rep. 34, 745–754 (2015).

6. Oltmanns, H. et al. Generation of backbone-free, low transgene copy plants by launching T-DNA from the Agrobacterium chromosome. Plant Physiol. 152, 1158–1166 (2010).

7. Ye, X. et al. Enhanced production of single copy backbone-free transgenic plants in multiple crop species using binary vectors with a pRi replication origin in Agrobacterium tumefaciens. Transgenic Res. 20, 773–786 (2011).

8. Szarzanowicz, M. J. et al. Binary vector copy number engineering improves Agrobacterium-mediated transformation. Nat. Biotechnol. 43, 1708–1716 (2025).

9. Francis, K. E. & Spiker, S. Identification of Arabidopsis thaliana transformants without selection reveals a high occurrence of silenced T-DNA integrations. Plant J. 41, 464–477 (2005).

10. Kim, S.-I., Veena & Gelvin, S. B. Genome-wide analysis of Agrobacterium T-DNA integration sites in the Arabidopsis genome generated under non-selective conditions. Plant J. 51, 779–791 (2007).

11. Ghedira, R., De Buck, S., Nolf, J. & Depicker, A. The efficiency of Arabidopsis thaliana floral dip transformation is determined not only by the Agrobacterium strain used but also by the physiology and the ecotype of the dipped plant. Mol. Plant Microbe Interact. 26, 823–832 (2013).

12. Napoli, C., Lemieux, C. & Jorgensen, R. Introduction of a chimeric chalcone synthase gene into Petunia results in reversible co-suppression of homologous genes *in trans*. Plant Cell 2, 279–289 (1990).

13. Głowacka, K. et al. An evaluation of new and established methods to determine T-DNA copy number and homozygosity in transgenic plants. Plant Cell Environ. 39, 908–917 (2016).

14. De Buck, S., Podevin, N., Nolf, J., Jacobs, A. & Depicker, A. The T-DNA integration pattern in Arabidopsis transformants is highly determined by the transformed target cell. Plant J. 60, 134–145 (2009).

15. Jupe, F. et al. The complex architecture and epigenomic impact of plant T-DNA insertions. PLoS Genet. 15, e1007819 (2019).

16. Pucker, B., Kleinbölting, N. & Weisshaar, B. Large scale genomic rearrangements in selected Arabidopsis thaliana T-DNA lines are caused by T-DNA insertion mutagenesis. BMC Genomics 22, 599 (2021).

17. Tinland, B. The integration of T-DNA into plant genomes. Trends Plant Sci. 1, 178–184 (1996).

18. Kleinboelting, N. et al. The Structural Features of Thousands of T-DNA Insertion Sites Are Consistent with a Double-Strand Break Repair-Based Insertion Mechanism. Mol. Plant 8, 1651–1664 (2015).

19. Thomson, G., Dickinson, L. & Jacob, Y. Genomic consequences associated with Agrobacterium-mediated transformation of plants. Plant J. 117, 342–363 (2024).

20. Nicolia, A., Ferradini, N., Veronesi, F. & Rosellini, D. An Insight into T-DNA Integration Events in Medicago sativa. Int. J. Mol. Sci. 18, (2017).

21. Olhoft, P. M., Flagel, L. E. & Somers, D. A. T-DNA locus structure in a large population of soybean plants transformed using the Agrobacterium-mediated cotyledonary-node method. Plant Biotechnol. J. 2, 289–300 (2004).

22. De Buck, S. T-DNA vector backbone sequences are frequently integrated into the genome of transgenic plants obtained by Agrobacterium-mediated transformation. Mol. Breed. 6, 459–468 (2000).

23. Afolabi, A. S., Worland, B., Snape, J. W. & Vain, P. A large-scale study of rice plants transformed with different T-DNAs provides new insights into locus composition and T-DNA linkage configurations. Theor. Appl. Genet. 109, 815–826 (2004).

24. Kuraya, Y. et al. Suppression of transfer of non-T-DNA vector backbone sequences by multiple left border repeats in vectors for transformation of higher plants mediated by Agrobacterium tumefaciens. Mol. Breeding 14, 309–320 (2004).

25. Gibson, D. G. et al. Enzymatic assembly of DNA molecules up to several hundred kilobases. Nat. Methods 6, 343–345 (2009).

26. Engler, C., Kandzia, R. & Marillonnet, S. A one pot, one step, precision cloning method with high throughput capability. PLoS ONE 3, e3647 (2008).

27. Kámán-Tóth, E., Pogány, M., Dankó, T., Szatmári, Á. & Bozsó, Z. A simplified and efficient Agrobacterium tumefaciens electroporation method. *3* Biotech 8, 148 (2018).

28. Clough, S. J. & Bent, A. F. Floral dip: a simplified method for *Agrobacterium*-mediated transformation of *Arabidopsis thaliana*. Plant J. 16, 735–743 (1998).

29. Rivero, L. et al. Handling *Arabidopsis* plants: growth, preservation of seeds, transformation, and genetic crosses. Methods Mol. Biol. 1062, 3–25 (2014).

30. Zhang, S. et al. Engineering *Rhodosporidium toruloides* for increased lipid production. Biotechnol. Bioeng. 113, 1056–1066 (2016).

31. de Groot, M. J., Bundock, P., Hooykaas, P. J. & Beijersbergen, A. G. *Agrobacterium tumefaciens*-mediated transformation of filamentous fungi. Nat. Biotechnol. 16, 839–842 (1998).

