## Supplemental Figures for "Binary vector origin predictably determines *Agrobacterium*-mediated transformation outcome across eukaryotic kingdoms"

**Affiliations:**

**Supplementary Figures**

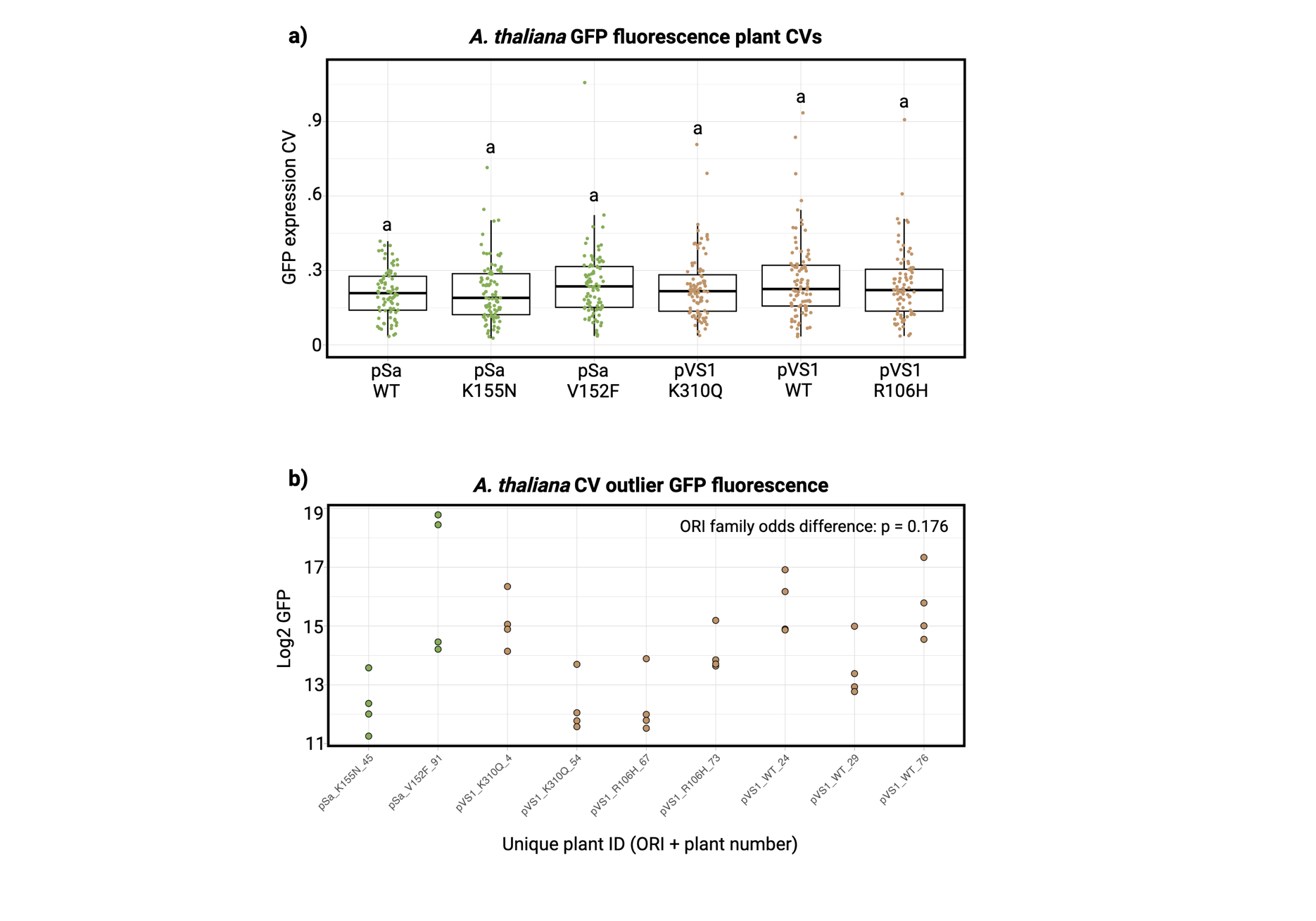

**Fig. S1** ***A. thaliana* intraplant GFP expression variance**

**a)** Boxplots of *A. thaliana* transformant CVs (standard deviation/mean) for GFP fluorescence taken from four leaves per transformant. Boxplots depict the interquartile range (IQR; Q1–Q3) with the median indicated by the central line. Whiskers extend to the most extreme values within 1.5 × IQR; points beyond are plotted as outliers. Distributions were compared by an ANOVA followed by a Tukey’s HSD test, with letters indicating significance groups (p > 0.05, n = 80-95 for each ORI variant group). Due to the observed leaf-to-leaf variation within samples, the mean GFP value was used for all further calculations conducted in this study. **b)** For outlier CV values above 0.6, log2 GFP fluorescence values were plotted for the four separate leaf measurements. Samples are shaded by ORI family and depict instances of high leaf-to-leaf variation in transgene expression. The incidence of high intra-plant expression variance was compared by ORI family (pSa: n = 260, pVS1: n = 267), and no significant enrichment was found (p = 0.176, Fisher’s Exact Test).

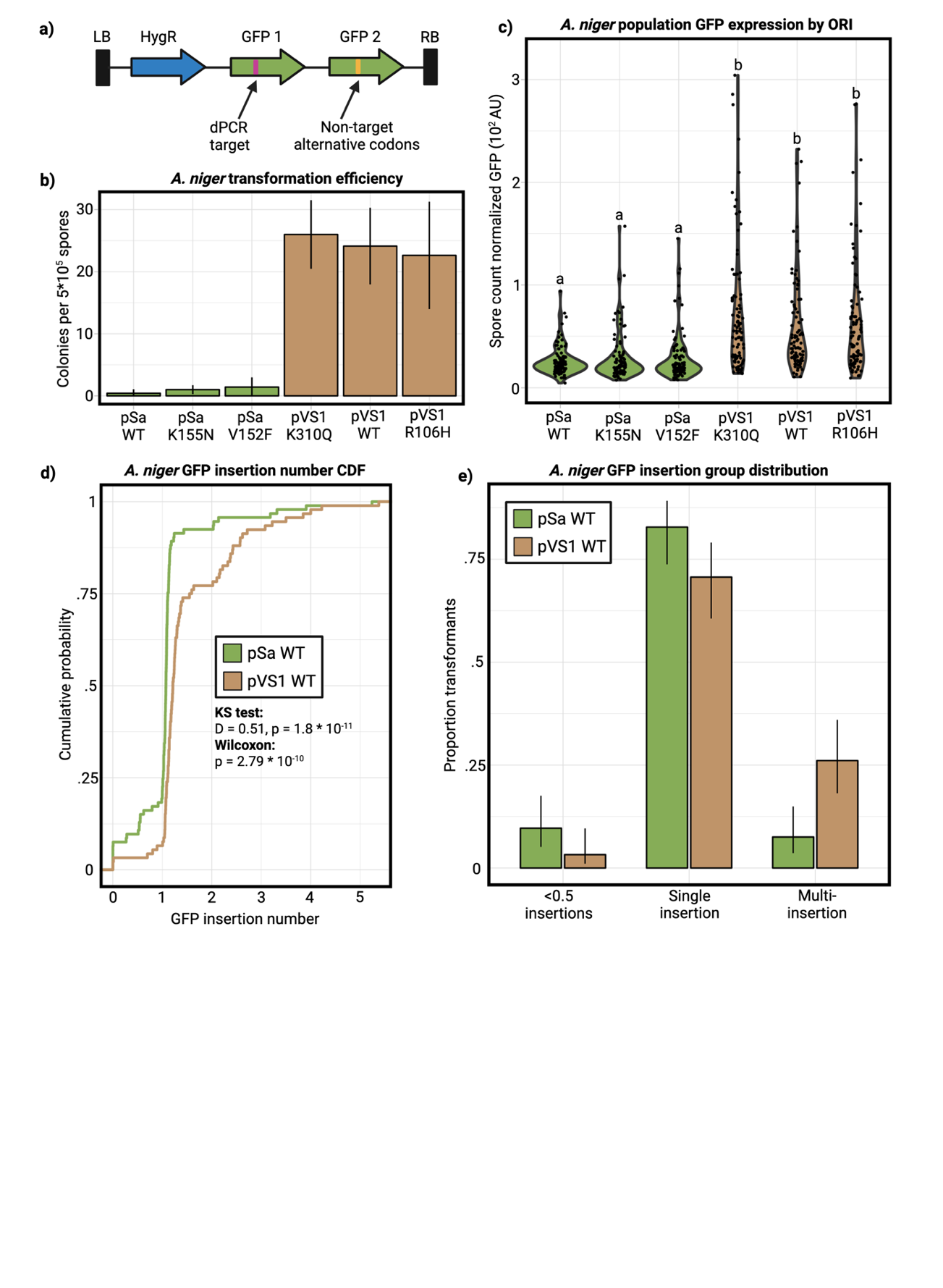

**Fig. S2 *Aspergillus niger* transformation overview**

**a)** Schematic of T-DNA for all *A. niger* binary vectors. This T-DNA contained two GFP cassettes to enable sufficient expression for GFP detection. One of these cassettes was used for dPCR quantification and the other was codon shuffled to not serve as a dPCR target. **b)** Transformation efficiency by ORI variant over four transformations. **c)** Distribution of GFP expression over 96 transformants by ORI variant. Expression distributions were compared by an ANOVA followed by a Tukey’s post hoc test (p < 0.05 for significance letters). **d)** GFP transgene insertion numbers were quantified with dPCR, and the population distributions for pSa WT and pVS1 WT are shown. Population differences were compared with a Kolmogorov-Smirnov test (D = 0.51, p = 1.8 * 10^-11^) and a Wilcoxon test (p = 2.79 * 10^-10^). **e)** Insertion outcomes were grouped (<0.5 insertions, single insertions, multi-insertions), and the estimated probabilities with Wald 95% confidence intervals were obtained from a binomial GLM followed by marginal means computed with emmeans.

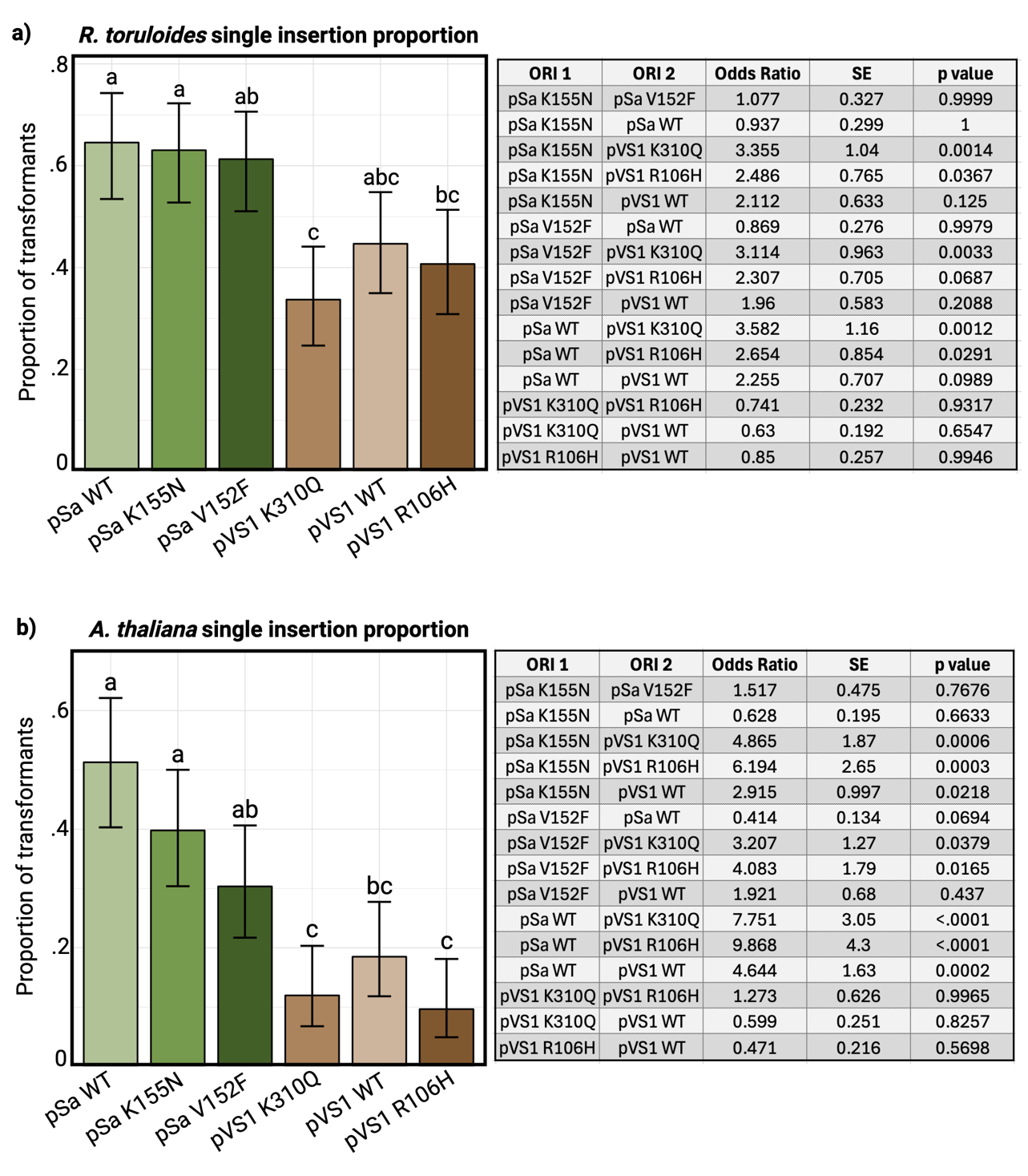

**Fig. S3: Single insertion rates by organism**

The single insertion rate for each ORI variant is shown for *R. toruloides* **(a)** and *A. thaliana* **(b)**. For each population (N = 79-94), the estimated marginal mean and 95% Wald CI derived from a binomial GLM are shown. Pairwise comparisons of each ORI variant were conducted, and letters above each bar denote Tukey significance groups (p < 0.05). Exact p-values are shown in the accompanying table.

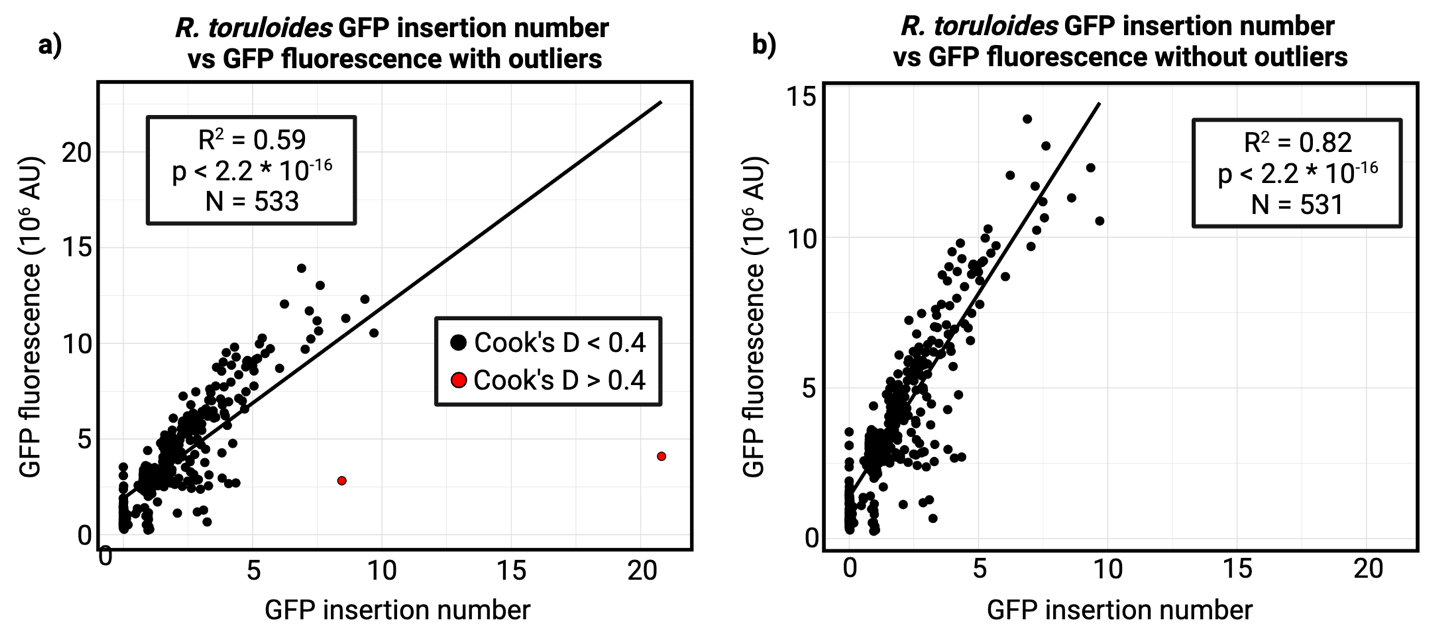

**Fig. S4: Identifying outliers in *R. toruloides* insertion number vs GFP expression regression**

Cook’s distance analysis of the R. *toruloides* dataset (N = 533) revealed two transformants with disproportionately high influence on the GFP–copy number regression (**a,** Cook’s D > 0.4; shown in red). To prevent these extreme outliers from dominating the regression outcome, these two points were excluded from subsequent analyses (**b**).

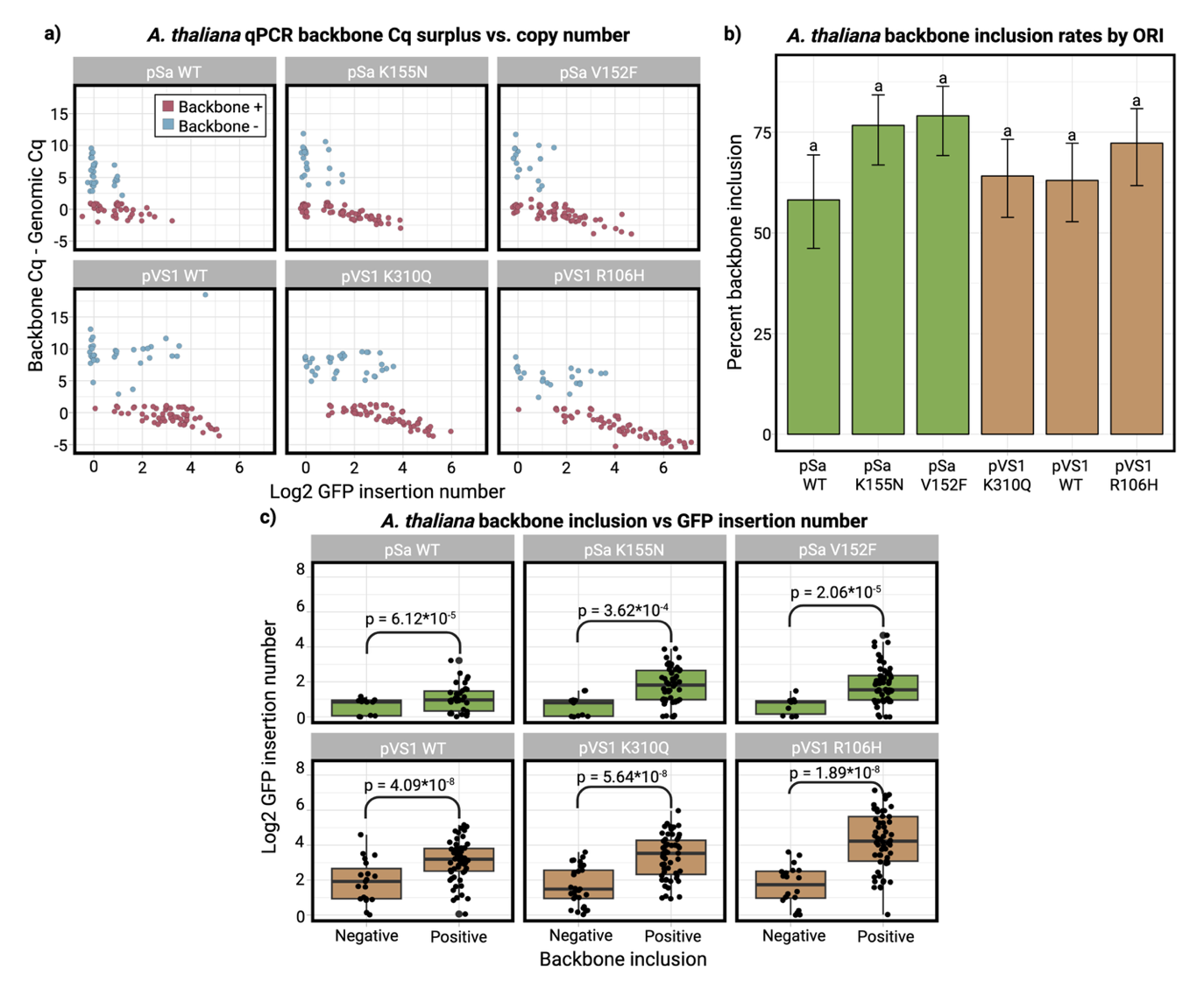

**Fig. S5: *A. thaliana* backbone analysis**

**a)** qPCR was used to analyze a backbone target and genomic target for each *A. thaliana* sample. The difference between the backbone target Cq and genomic target Cq (backbone target Cq surplus) was used to classify backbone inclusion. A positive backbone target Cq surplus indicates less backbone target template within the sample compared to the genomic target template, with each N integer value corresponding to ~$(\frac{1}{2})^{N}$ the template number. Surplus backbone Cq values over 2 were classified as backbone negative (blue) and clustered well from <2 backbone Cq surplus samples (red). Surplus backbone Cq is plotted against log2 GFP insertions, showing clustering. **b)** The rate of backbone inclusion by ORI variant was calculated using a binomial GLM-derived estimated marginal mean along with a 95% Wald CI. A Holm-corrected comparison found no significant difference in inclusion rate between ORI variants (p > 0.05). **c)** Backbone negative samples had lower median GFP insertion numbers compared to backbone positive samples for all ORI variants (Wilcoxon test, p < 0.05).

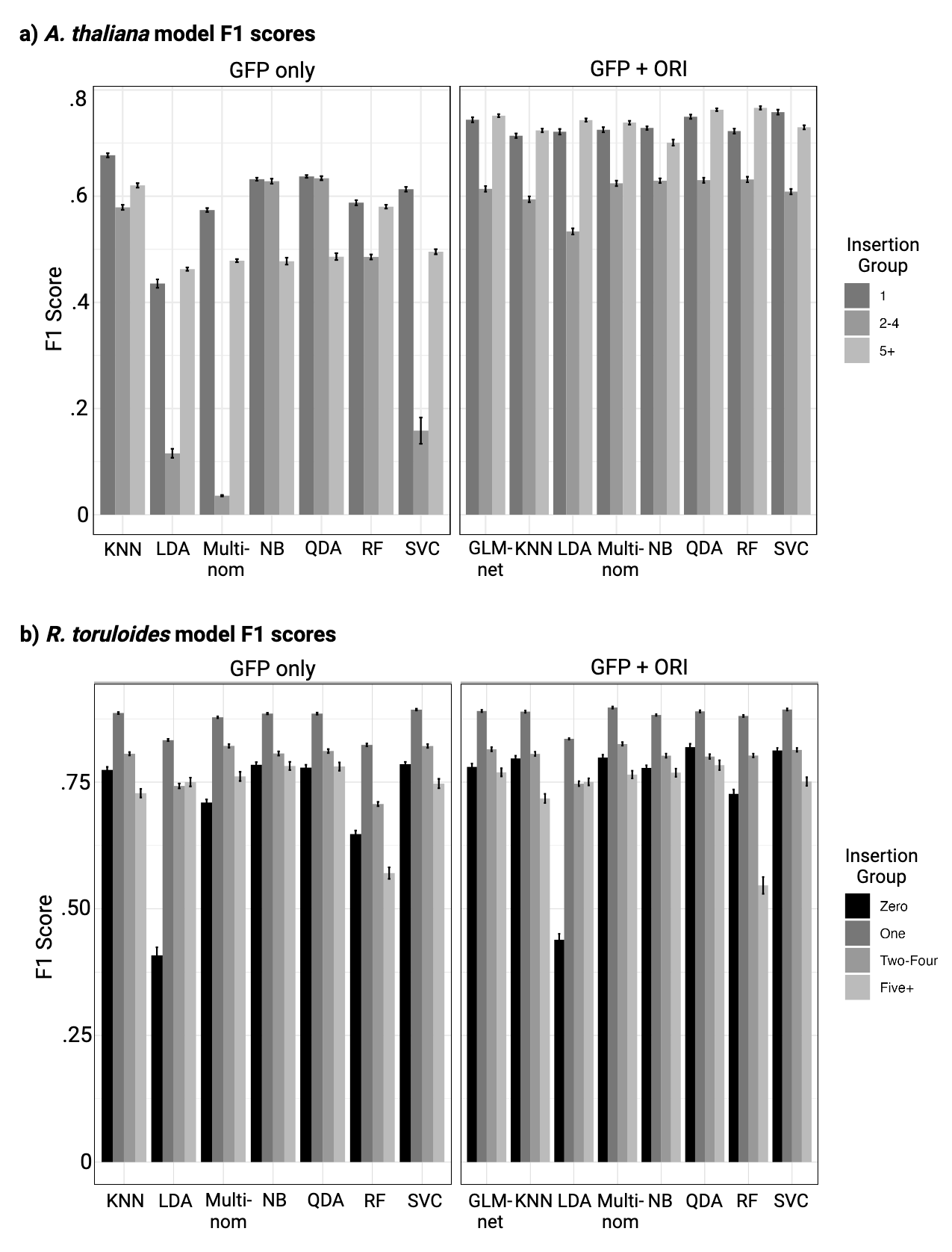

**Fig. S6: Model performance for predicting transgene insertion number classes**

**(a)** A. thaliana and **(b**) R. toruloides. Bars show mean F1 scores for each insertion number class predicted from either GFP alone or GFP plus ORI family, across eight supervised learning models: quadratic discriminant analysis (QDA), linear discriminant analysis (LDA), k-nearest neighbors, random forest (RF), multinomial logistic regression (multinom), support vector classifier (SVC; linear kernel), naïve Bayes (NB), and penalized multinomial regression with elastic net regularization (glmnet). For each model, data were randomly partitioned into 70% training and 30% withheld test sets, repeated across 100 iterations. Error bars represent the standard error of the mean (SEM) of F1 scores across these 100 train/test replicates.

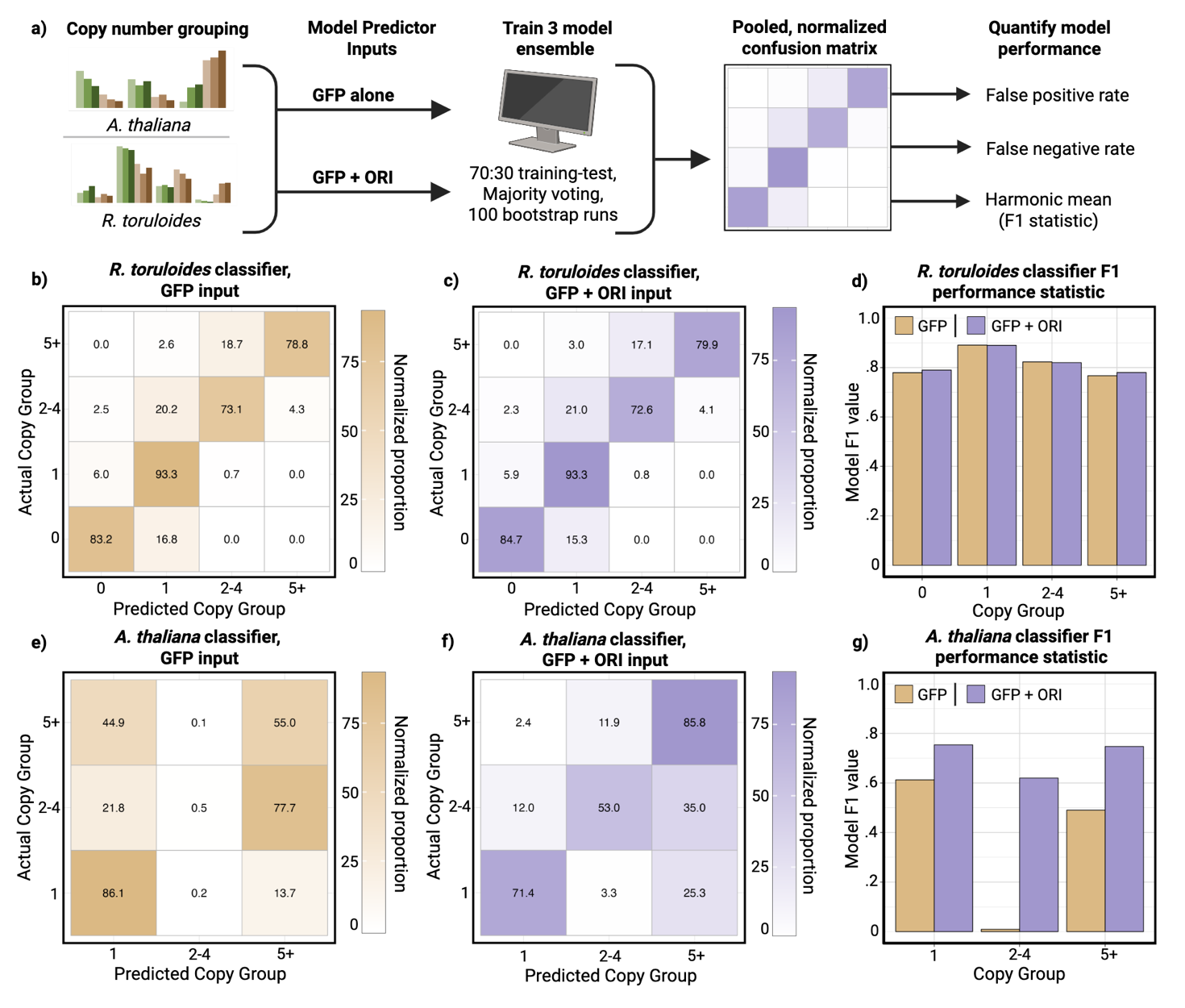

**Fig. S7: Expanded ensemble performance of GFP insertion number**

An expansion of Fig. 5. **(a)** An ensemble of predictive models was trained to predict the GFP insertion number of a transformant based on either GFP output alone or GFP + ORI identity as predictive factors. These data were used to conduct 100 bootstrapped 70:30 training-test splits across three independent models (SVC, NB, and QDA for *A. thaliana* and SVC, NB, and multinomial logistic regression model for *R. toruloides*), which were then combined in a majority voting ensemble. Confusion matrices were generated from the bootstrapped runs to analyze the ensemble’s performance and rates of false positive and negative results for *R. toruloides* (**b, c**) and *A. thaliana* (**e, f**). The ensemble F1 statistics per insertion group are reported for *R. toruloides* in **(d)** and *A. thaliana* in **(g)**.

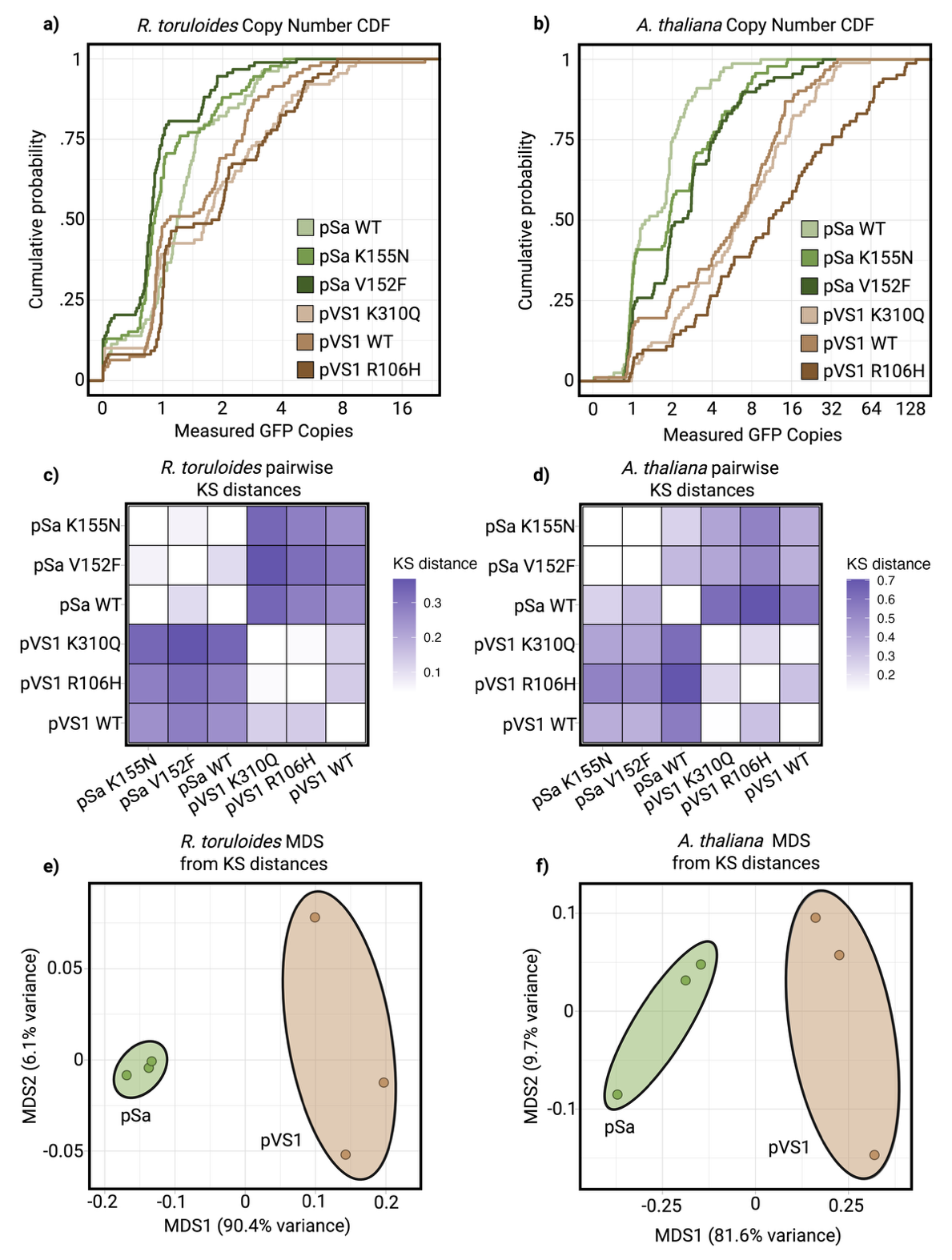

**Fig. S8: Multi-dimensional scaling analysis of *R. toruloides* and *A. thaliana***

CDFs for GFP insertion number in *R. toruloides* **(a)** and *A. thaliana* **(b)**. Using these CDFs, the pairwise KS distances between each ORI variant population can be calculated. These KS distances are displayed in **(c)** and **(d)** for each organism. Using these matrices, MDS plots can be generated for each organism **(e)** and **(f)**, with the percent variance for each axis displayed accordingly. For both organisms, the vast majority of the variance is found on the x axis, separating the six variants by ORI family.

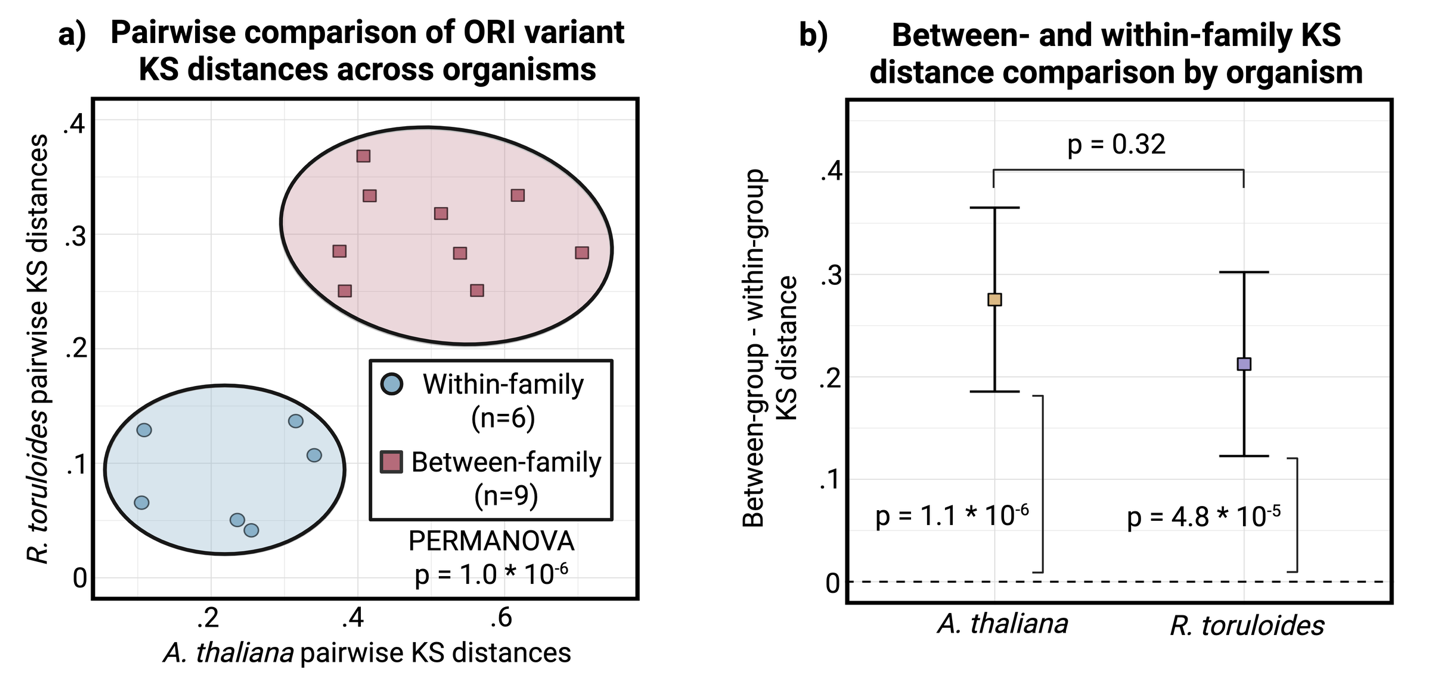

**Figure S9: Differences in within- vs. between-family KS statistics by organism**

**a)** Pairwise KS distances for all ORI variants were plotted by organism. Points across organisms significantly clustered for within-family (n = 6) and between-family (n = 9) points (PERMANOVA, p = 1.0*10^-6^). **b)** For each organism, the plotted square shows the estimated marginal mean difference between *between-family* and *within-family* KS distances (between − within), with 95% Wald confidence intervals. Both organisms exhibit values significantly greater than zero –shown as a dotted line at y = 0—(Arabidopsis: p = 1.1 × 10⁻⁶; *R. toruloides*: p = 4.8 × 10⁻⁵), indicating stronger divergence across ORI families than within families. The magnitude of this difference does not significantly differ between organisms (p = 0.32).
